# Universal patterns of selection in cancer and somatic tissues

**DOI:** 10.1101/132324

**Authors:** Iñigo Martincorena, Keiran M. Raine, Moritz Gerstung, Kevin J. Dawson, Kerstin Haase, Peter Van Loo, Helen Davies, Michael R. Stratton, Peter J. Campbell

**Author notes:** Correspondence: I.M., P.J.C.

## Abstract

Cancer develops as a result of somatic mutation and clonal selection, but quantitative measures of selection in cancer evolution are lacking. We applied methods from evolutionary genomics to 7,664 human cancers across 29 tumor types. Unlike species evolution, positive selection outweighs negative selection during cancer development. On average, <1 coding base substitution/tumor is lost through negative selection, with purifying selection only detected for truncating mutations in essential genes in haploid regions. This allows exome-wide enumeration of all driver mutations, including outside known cancer genes. On average, tumors carry ∼4 coding substitutions under positive selection, ranging from <1/tumor in thyroid and testicular cancers to >10/tumor in endometrial and colorectal cancers. Half of driver substitutions occur in yet-to-be-discovered cancer genes. With increasing mutation burden, numbers of driver mutations increase, but not linearly. We identify novel cancer genes and show that genes vary extensively in what proportion of mutations are drivers versus passengers.

**HIGHLIGHTS:**

- Unlike the germline, somatic cells evolve predominantly by positive selection
- Nearly all (∼99%) coding mutations are tolerated and escape negative selection
- First exome-wide estimates of the total number of driver coding mutations per tumor
- 1-10 coding driver mutations per tumor; half occurring outside known cancer genes

## INTRODUCTION

Somatic cells accumulate mutations throughout life. These mutations can be classified into those that confer a selective advantage on the cell, increasing survival or proliferation, (so-called ‘driver’ mutations); those that are selectively neutral; and those that are disadvantageous to the cell and result in its death or senescence (Stratton et al., 2009). Cancer is one end-product of mutation in somatic cells, in which a single clonal lineage acquires a complement of driver mutations that enables the cells to evade normal constraints on cell proliferation, invade tissues, and spread to other organs.

While the general principles of cancer evolution have been well documented for some decades (Cairns, 1975; Nowell, 1976; Stratton et al., 2009), a number of fundamental questions remain unanswered. We still do not have accurate estimates of the number of mutations required to drive a cancer, and whether this varies extensively across tumor types or with different mutation rates (Martincorena and Campbell, 2015). One approach to this question has been to use age-incidence curves to estimate the number of rate-limiting steps required for a cancer to develop (Armitage and Doll, 1954; Tomasetti et al., 2015), with the implicit assumption of a one-to-one correspondence between rate-limiting steps and driver mutations. However, not all driver mutations need be rate-limiting (Yates et al., 2015), nor every rate-limiting event need be a driver mutation (Martincorena and Campbell, 2015). A second approach to estimating the number of driver mutations has simply been to count the mutations occurring in known cancer genes, but this is limited by incomplete lists of cancer driver genes and by the presence of passenger mutations in cancer genes. Thus, despite its fundamental importance, the issue of how many somatic mutations drive a cancer remains unresolved.

A second major gap in our understanding of cancer evolution is that we have not yet been able to measure the importance of negative selection in shaping the cancer genome, and to what extent somatic lineages expire due to the effects of deleterious mutations. Detection of negative selection in cancer genomes is a potentially important endeavor as it may help identify genes essential for cancer growth and patterns of synthetic lethality, potentially yielding a novel class of therapeutic targets. Also, with increasing interest in the role of neoantigens created by somatic mutations shaping the immune response to cancer (McGranahan et al., 2016; Rajasagi et al., 2014; Rooney et al., 2015), we might expect that purifying selection would suppress clones with mutations that elicit strong immune reaction.

Thirdly, while we have increasingly detailed lists of cancer genes (Kandoth et al., 2013; Lawrence et al., 2014; Vogelstein et al., 2013), it is not always straightforward to identify which mutations in those genes are true driver mutations nor how many mutations in other genes might be drivers. This will become an increasingly important question as cancer genome sequencing moves into routine clinical practice – therapeutic decision support for an individual patient critically depends on accurate identification of which specific mutations drive the biology of that person’s cancer (Gerstung et al., 2017).

In this study, we address these three open questions by adapting methods from evolutionary genomics to the study of cancer genomes. The key advance in the models we develop is that we can directly enumerate the excess or deficit of mutations in a given gene, a group of genes or even at whole-exome level, compared to the expectation for the background mutational processes. This enables us to provide robust estimates of the total number of coding driver mutations across cancers; how many coding point mutations are lost through negative selection and a detailed dissection of the distribution of driver mutations in individual cancer genes across different tumor types.

## RESULTS

### Quantitative assessment of positive and negative selection

Detection of selection in traditional comparative genomics typically requires a measure of the expected density of selectively neutral mutations in a gene. In the context of cancer, a gene under positive selection will carry an extra complement of driver mutations in addition to neutral (passenger) mutations – it is this recurrence of mutations across cancer patients that has underpinned discoveries of cancer genes from the Philadelphia chromosome to modern genomic studies (Martincorena and Campbell, 2015). A gene subject to purifying selection of deleterious mutations would have fewer mutations than expected under neutrality (Greenman et al., 2006).

Building on previous work (Greenman et al., 2006; Martincorena et al., 2015; Yang et al., 2003), we use dN/dS, the ratio of non-synonymous to synonymous mutations, to quantify selection in cancer genomes. This relies on the assumption that the vast majority of synonymous mutations are selectively neutral and hence a good proxy to model the expected mutation density (we address the accuracy of this assumption later; Supplementary Methods S5.3). dN/dS has a long history in the study of selection in species evolution (Goldman and Yang, 1994; Nei and Gojobori, 1986; Yang and Bielawski, 2000), but several modifications are required for somatic evolution.

The first critical refinement is more comprehensive models for context-dependent mutational processes (Alexandrov et al., 2013; Greenman et al., 2006; Yang et al., 2003). Traditional implementations of dN/dS use simplistic mutation models that lead to systematic bias in dN/dS ratios, and can cause incorrect inference of positive and negative selection (**Fig. S1A**) – such biases have affected previous studies in this area (Ostrow et al., 2014) (Supplementary Text S6). We therefore use a model with 192 rate parameters that accounts for all 6 types of base substitution, all 16 combinations of the bases immediately 5’ and 3’ to the mutated base and transcribed versus non-transcribed strands of the gene (Supplementary Methods S1). A second refinement is the addition of other types of non-synonymous mutations beyond missense mutations, including nonsense and essential splice site mutations (Greenman et al., 2006), and a method for small insertions and deletions (indels). Thirdly, extreme caution was exercised during variant calling to avoid biases emerging from germline variants, since these have a much lower dN/dS ratio than somatic mutations (**Fig. 1**) (Supplementary Methods S2). Misannotation of a germline polymorphism as a somatic mutation will bias somatic dN/dS downwards; excessively filtering true somatic mutations that occur at positions known to be polymorphic in the population will bias somatic dN/dS upwards (**Fig. S1B**). For example, we can demonstrate that germline contamination of the public mutation catalogs from several datasets in The Cancer Genome Atlas is responsible for a false signal of negative selection in them (Supplementary Text S8, **Fig. S1C**). Fourthly, to detect selection at the level of individual genes reliably, and particularly for driver gene discovery, we refined dN/dS to consider the variation of the mutation rate along the human genome. A simple way to do so is estimating a separate mutation rate for every gene (Wong et al., 2014), but this approach has poor statistical efficiency with current sample sizes. Instead, we developed a statistical model (*dNdScv*) that combines the local observed synonymous mutation rate with a regression model using covariates that predict the variable mutation rate across the genome (Lawrence et al., 2013; Polak et al., 2015; Schuster-Bockler and Lehner, 2012) (Supplementary Methods S1.3 and Supplementary Text S9). This approach has the advantage of optimizing the balance between local and global data on estimating background mutation rates to provide a statistically efficient inference framework for departures from neutrality (**Fig. S1F,G**).

**Fig. 1.**
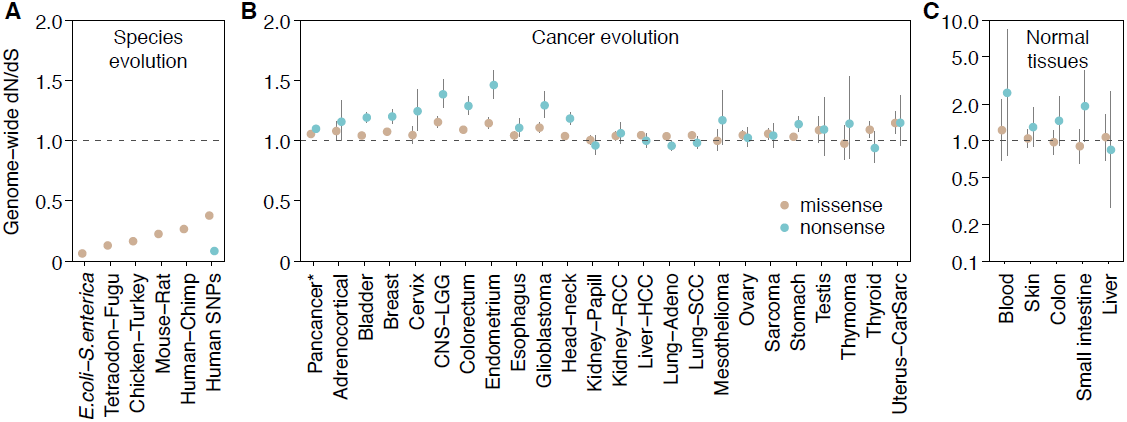
Genome-wide dN/dS ratios show a distinct pattern of selection universally shared across cancer types. (**A**) Species evolution: median dN/dS ratios across genes for missense mutations (data from (Martincorena et al., 2012), Ensembl (Vilella et al., 2009)). Data on germline human SNPs are from the 1000 genomes phase 3 (Auton et al., 2015), restricted to SNPs with minor allele frequency >=5%. (**B**) Cancer evolution: genome-wide dN/dS values for missense and nonsense mutations across 23 cancer types (Supplementary Methods S2.3). (**C**) Somatic mutations in normal tissues (data from (Blokzijl et al., 2016; Martincorena et al., 2015; Welch et al., 2012)). Error bars depict 95% confidence intervals.

In order to study the landscape of positive and negative selection in cancer, we applied these approaches to a collection of 7,664 tumors from 29 cancer types from The Cancer Genome Atlas (Table S1). Somatic mutations were re-called with our in-house algorithms across 24 cancer types to ensure comparability across tumor types and avoid biases from germline polymorphisms (Supplementary Methods S2).

### A universal and distinct pattern of selection in cancer

Comparative genomic studies of related species typically reveal very low dN/dS ratios, reflecting that the majority of germline non-synonymous mutations are removed by negative selection over the course of evolution. For example, comparison of orthologous genes from *Escherichia coli* and *Salmonella enterica* yields an average dN/dS∼0.06 across genes. This indicates that at least ∼94% of missense mutations have been removed by negative selection. The dN/dS ratio for nonsense mutations in common human germline polymorphisms is similarly low (dN/dS∼0.08). dN/dS ratios vary across species but a pattern of overwhelming negative selection invariably characterizes species evolution (**Fig. 1A**).

In stark contrast, cancer evolution shows a pattern in which dN/dS ratios are close to, but slightly above, one (**Fig. 1B**). This pattern is universally shared across tumor types studied here and applies to both missense and truncating substitutions (nonsense and essential splice site mutations). This indicates that mutations under positive selective pressure are somewhat more numerous in cancers than mutations under negative selection, but the overall picture is close to neutrality. Importantly, similar values of dN/dS around or above one are found in somatic mutations detected in healthy tissues, including blood, skin, liver, colon and small intestine (Blokzijl et al., 2016; Martincorena et al., 2015; Welch et al., 2012) (**Fig. 1C**). Although these data are still limited, dN/dS∼1 appears to characterize somatic evolution in normal somatic tissues as well as all cancers that we have studied so far.

### Identification of genes under positive selection

By definition, driver genes are genes under positive selection in cancer genomes. To show the ability of dN/dS to uncover driver genes, we used *dNdScv* to identify genes for which dN/dS was significantly higher than 1 (Supplementary Methods S3), both across all 7,664 cancers and for each tumor type individually (**Fig. 2A**). This revealed 179 cancer genes under positive selection at 5% false discovery rate. Of these, 54% are canonical cancer genes present in the Cancer Gene Census (Forbes et al., 2015). Using restricted hypothesis testing (Lawrence et al., 2014) on *a priori* known cancer genes identifies an additional 24 driver genes (Supplementary Methods S1.4). Evaluation of genes not present in the Census reveals that most have been previously reported as cancer genes, have clear links to cancer biology or have been found in other pan-cancer analyses (Kandoth et al., 2013; Lawrence et al., 2014; Rubio-Perez et al., 2015) (Table S2). Novel candidate cancer genes include *ZFP36L1* and *ZFP36L2*, which have recently been shown to promote cellular quiescence and suppress S-phase transition during B cell development (Galloway et al., 2016). We find higher than expected rates of inactivating mutations in the two genes in several tumor types, suggesting that they have a tumor suppression role. Other novel tumor suppressor genes identified here include *KANSL1*, a scaffold protein for histone acetylation complexes (Dias et al., 2014); *BMPR2*, a receptor serine/threonine kinase for bone morphogenetic proteins; *MAP2K7*, involved in MAP-kinase signaling; and *NIPBL*, a member of the cohesin complex.

**Fig. 2.**
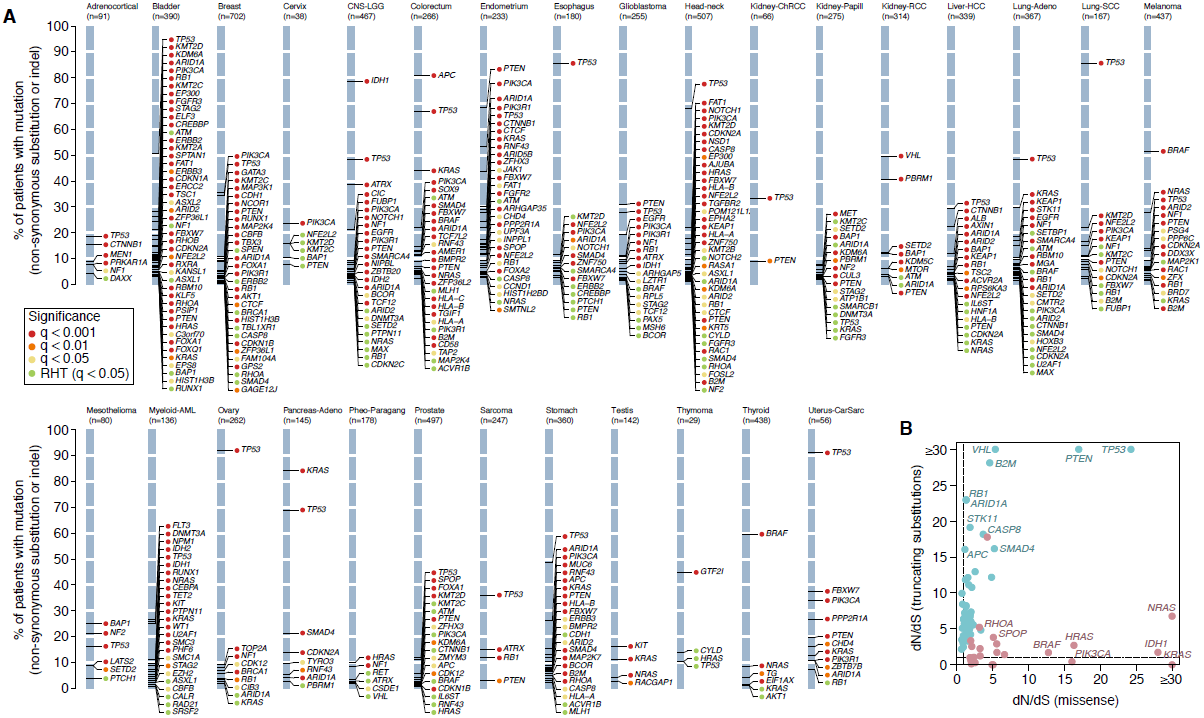
Positively selected genes in cancer genomes. (**A**) List of genes detected under significant positive selection (dN/dS>1) in each of the 29 cancer types. Y-axes show the percentage of patients carrying a non-synonymous substitution or an indel in each gene. The color of the dot reflects the significance of each gene. RHT = Restricted Hypothesis Testing on known cancer genes (Supplementary Methods S3 and Table S2). (**B**) Pancancer dN/dS values for missense and nonsense mutations for genes with significant positive selection on missense mutations (depicted in red) and/or truncating substitutions.

As expected, depending on whether nonsense or missense mutations predominate, genes generally fall into two classes: oncogenes, with strong selection on missense mutations, or tumor suppressor genes, with stronger selection on truncating mutations (**Fig. 2B**). Significant dN/dS ratios reach very high values in frequently mutated driver genes, often higher than 10 or even 100 (**Fig. 2B**). This gives quantitative information about the proportion of driver mutations. For example, dN/dS=10 for a gene evidences that there are ten times more non-synonymous mutations in the gene than expected under neutral accumulation of mutations, indicating that at least ∼90% of the non-synonymous mutations in the gene are genuine driver mutations (Greenman et al., 2006).

### Negative selection is largely absent for coding substitutions

While some somatic mutations can confer a growth advantage, others may impair cell survival or proliferation. Clones carrying such mutations would senesce or die, with the result that the mutation would be lost from the catalog of variants seen in the eventual cancer. This negative or purifying selection will lead to dN/dS<1 in a given gene or set of genes if it occurs at appreciable rates. Negative selection on somatic mutations has been long anticipated (Beckman and Loeb, 2005; McFarland et al., 2014; Nowell, 1976; Stratton et al., 2009), but not yet reliably documented in cancer genomes. This is due to the fact that statistical detection of lower mutation density than expected by chance requires large datasets and very careful consideration of mutation biases and germline SNP contamination. Previous studies have reported extensive negative selection in cancer genomes (Ostrow et al., 2014), but we find that this conclusion is erroneous, resulting from overly simplistic and biased mutation models (Supplementary Text S6 and **Fig. S1A**). Further, even just a few percent of germline polymorphisms contaminating catalogs of somatic mutations can lead to false signals of negative selection (**Fig. S1B**), a problem that we detect in the public mutation calls of several TCGA datasets (Supplementary Text S8 and **Fig. S1C**). Owing to these limitations, the true extent of negative selection in cancer evolution remains unclear.

To determine the potential extent of negative selection, we first studied the distribution of observed dN/dS values per gene. There is considerable spread of these observed values around the neutral peak at dN/dS=1.0 (**Fig. 3A**), which at face value might suggest that many genes are under positive or negative selection. However, the limited numbers of mutations per gene make individual dN/dS values noisy, and we find that the observed distribution almost exactly matches that seen in simulations under a model where all genes are neutral (Supplementary Methods S4.2.1). To formally estimate the fraction of genes under negative selection, we infer the underlying distribution of dN/dS values from the observed data using a binomial mixture model (Supplementary Methods S4.2.2) (**Fig. 3B-C**). We find that the vast majority of genes are expected to accumulate point mutations near neutrally, with dN/dS∼1. A small fraction of genes (∼2.2%; CI_95%_=1.0-3.9%) show dN/dS≥1.5, consistent with current estimates of the numbers of cancer genes. Only a tiny fraction of genes (∼0.14%; CI_95%_=0.02-0.51%), approximately a few tens, are estimated to exhibit negative selection with dN/dS≤0.75 (**Fig. 3C**; **Fig. S2**).

**Fig. 3.**
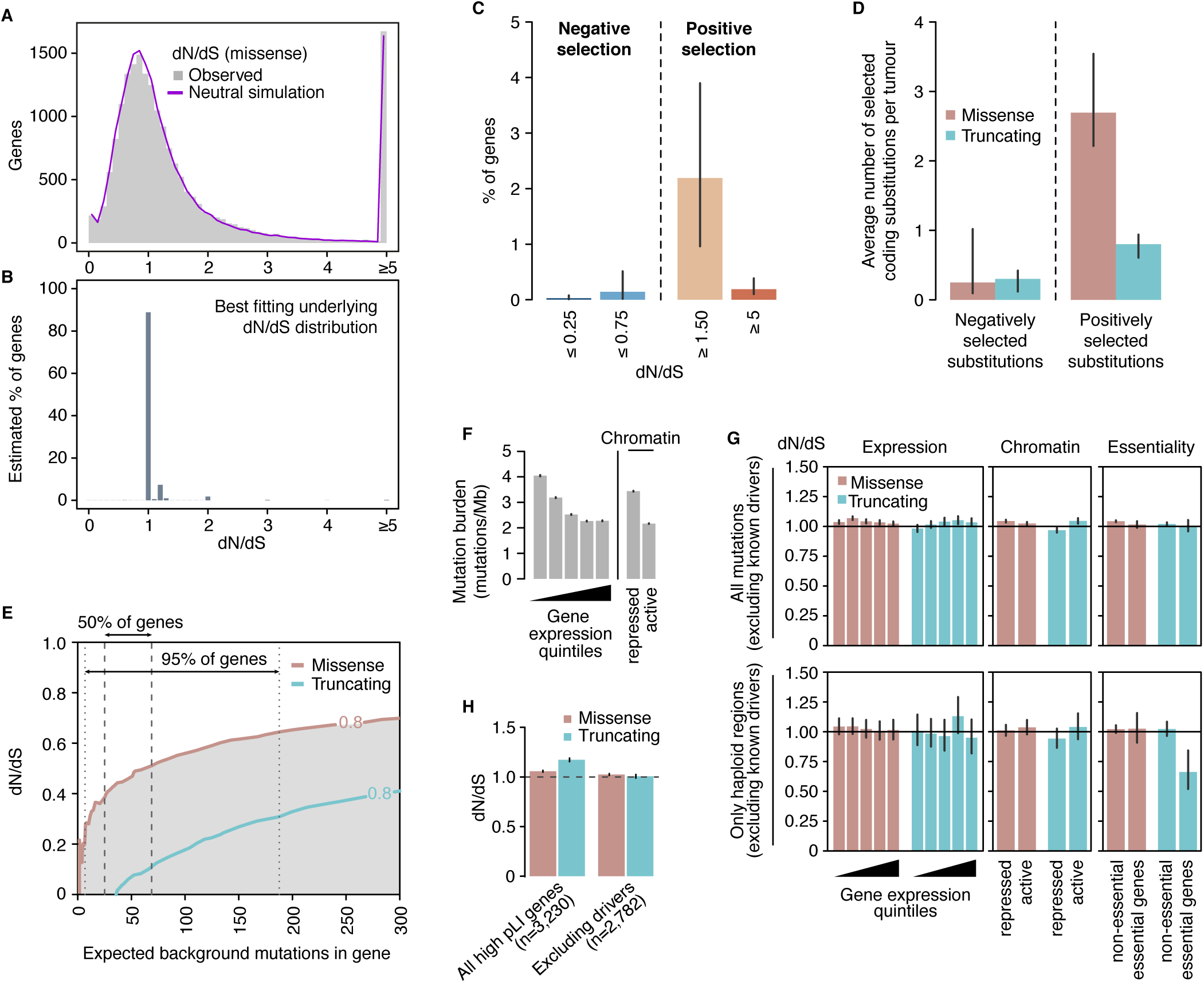
Negative selection in cancer. (**A**) Distributions of dN/dS values per gene for missense mutations in non-LOH regions (Supplementary Methods S4.1). The real distribution is shown in grey and the distribution observed in a neutral simulation is shown in purple (Supplementary Methods S4.2.1). (**B**) Underlying distribution of dN/dS values across genes inferred from the observed distribution (Supplementary Methods S4.2.2). (**C**) Estimated percentage of genes under different levels of positive and negative selection based on the inferred dN/dS distribution in **Fig. 3B**. (**D**) Average number of selected mutations per tumor based on the inferred distributions of dN/dS across genes, combining missense and truncating mutations from all copy number regions (Supplementary Methods S4.2.3). Error bars depict 95% confidence intervals. (**E**) Power calculation for the statistical detection of negative selection (dN/dS<1) as a function of the extent of selection (dN/dS) and the neutrally-expected number of mutations in a gene in a cohort (Supplementary Methods S4.3). Shaded areas under the curves reflect power>80%. Vertical lines indicate the range in which the middle 50% and 95% of genes are, in the dataset of 7,664 tumors. (**F**) Average mutation burden in genes grouped according to gene expression quintile and chromatin state. (**G**) Average dN/dS values for genes grouped according to gene expression quintile, chromatin state and essentiality (Supplementary Methods S4.4). (**H**) Average dN/dS values for all mutations in genes found to be haploinsufficient in the human germline, including and excluding putative driver genes. Haploinsufficient genes are defined as those having a pLI score >0.9 in the ExAC database (Lek et al., 2016).

These distributions also enable us to obtain approximate estimates of the average number of coding substitutions lost by negative selection per tumor (**Fig. 3D**; Supplementary Methods S4.2.3). On average across this diverse collection of tumors, less than one coding substitution per tumor (0.55/patient; CI_95%_=0.31-1.16) appears to have been lost by negative selection, accounting for <1% of all coding mutations. We note the formal possibility that dN/dS=1 can occur when the numbers of positively and negatively selected mutations in a given gene are exactly balanced. This could lead us to underestimate the extent of negative selection, but only if a large number of genes showed such an exact balance, which appears very unlikely.

Although negative selection in cancers might be weak globally, it remains possible that negative selection may act in very specific scenarios, genes or gene sets. No single gene had a dN/dS significantly less than 1 after multiple hypothesis testing correction, even if we boost our power by performing restricted hypothesis testing on 1,734 genes identified by *in vitro* screens as essential (Blomen et al., 2015) (Supplementary Methods S4.3). To address the possibility of making a type II inference error, we evaluated our statistical power to detect negative selection at the level of individual genes in this dataset (**Fig. 3E**) (Supplementary Methods S4.3). We found that there is enough power to detect negative selection at dN/dS<0.5 on missense mutations for most genes in the dataset, but we have less power for detecting negative selection acting on truncating mutations (**Fig. 3E**). Thus, the lack of significant negative selection in any gene in the current dataset reveals that negative selection is weaker than these detection limits.

We next examined whether specific groups of genes might be subject to negative selection, after excluding 987 putative cancer genes to avoid obscuring the signal of negative selection (Supplementary Methods S4.4). Sets of genes that may be expected to be under stronger negative selection include highly expressed genes or genes in active chromatin regions. Lower mutation density has been observed in cancer genomes in highly expressed genes and open chromatin (**Fig. 3F**) (Pleasance et al., 2010b; Schuster-Bockler and Lehner, 2012), and some have suggested that this may be a signal of negative selection (Lee et al., 2010). However, we found that dN/dS values are virtually indistinguishable from neutrality for both missense and truncating substitutions across gene expression levels and chromatin states. This confirms that the lower density of mutations observed in open chromatin and highly expressed genes is due to lower mutation rates in these regions and not negative selection. The lack of detectable negative selection even extends to nonsense mutations in essential genes (**Fig. 3G**; top panel). Gene sets grouped by gene ontology and functional annotation similarly revealed no clear evidence of negative selection (**Fig. S3**).

One reason for this unexpected weakness of negative selection in cancer could be that cancer cells typically carry two (or more) copies of most genes, reducing the impact of mutations inactivating a single gene copy. We used copy number data for the samples studied here to identify those coding mutations occurring in haploid regions of the genome. Strikingly, most missense and even truncating substitutions affecting the single remaining copy of a gene seem to accumulate at a near-neutral rate, suggesting that they are largely tolerated by cancer cells (**Fig. 3G**; bottom panel). However, for essential genes in regions of copy number 1, nonsense substitutions do exhibit significantly reduced dN/dS, with about one third of such variants lost through negative selection (dN/dS=0.66, *P*-value=8.4×10^-4^, **Fig. 3G**). Analysis of mutations in haploinsufficient genes in human evolution also revealed no detectable negative selection in cancer evolution, either on missense or truncating substitutions (**Fig. 3H**).

Overall, these analyses show that negative selection in cancer genomes is much weaker than anticipated. With the exception of driver mutations, nearly all coding substitutions (∼99%) appear to accumulate neutrally during cancer evolution and are tolerated by cancer cells. Several factors are likely to contribute to the weakness of negative selection, including the buffering effects of two or more copies of most genes and the fact that many genes are presumably dispensable to somatic cells (Morley, 1995). Further, in cancer, the regular occurrence of strong driver mutations generating selective sweeps will enable weakly deleterious mutations not yet expunged to hitchhike on the clonal expansion, reducing the efficiency of negative selection to remove deleterious variants (McFarland et al., 2013) (see Supplementary Text S10 for a discussion on factors limiting negative selection in somatic evolution).

Immune surveillance is believed to be a relevant force shaping cancer evolution, potentially acting to purge clones carrying neoantigens generated by somatic mutations. Genomic studies have predicted that cancers typically carry tens of coding mutations that generate potential neoantigens (McGranahan et al., 2016; Rajasagi et al., 2014), with as many as 50% of non-synonymous mutations predicted to create a neoantigen (Rooney et al., 2015). The observation that ∼99% of somatic point mutations are tolerated and accumulate neutrally in cancer cells confirms that the vast majority of predicted neoantigens do not elicit an immune response capable of eradicating the clone in normal conditions, even if they could be exploited therapeutically (Stronen et al., 2016).

### Number of driver mutations per tumor

The number of driver mutations required to generate a tumor has been a long-standing question in cancer (Armitage and Doll, 1954; Martincorena and Campbell, 2015; Nordling, 1953; Stratton et al., 2009; Tomasetti et al., 2015). Early epidemiological observations using age-incidence statistics suggested that cancer may result from a small number of rate-limiting steps, around five to seven, possibly driver mutations (Armitage and Doll, 1954; Nordling, 1953). The sequencing of thousands of cancer genomes has not clarified this question further since it remains unclear what fraction of non-synonymous mutations observed in known cancer genes are genuine positively-selected driver mutations and, particularly, how many driver mutations occur in cancer genes that are yet to be discovered. This has recently led some to propose that a large number of the mutations seen in any given tumor may have been positively selected, lowly recurrent across patients but playing a driver role in the specific genomic context of that patient (Castro-Giner et al., 2015).

From an observed dN/dS ratio, we can estimate the number of extra non-synonymous mutations over what would have been expected under neutrality (Greenman et al., 2006) (Supplementary Methods S5). Since we have shown that negative selection does not significantly deplete non-synonymous mutations in somatic evolution, this additional burden is a direct estimate of the number of driver substitutions. For example, combining all coding mutations observed in 369 cancer genes across 689 breast cancer samples yields dN/dS=1.95 (CI_95%_: 1.72-2.21). This implies that there are 1.95x more non-synonymous mutations than expected neutrally, or, equivalently, 49% (CI_95%_: 42%-55%) of the observed non-synonymous mutations are positively selected driver mutations (**Fig. 4A**). Although this calculation does not inform which of these mutations are drivers, it provides a statistical framework for inferring the fraction and the absolute number of drivers in a catalog of mutations. Interestingly, manual annotation of breast cancer genomes has led to very similar estimates of the number of driver mutations in known cancer genes per tumor (**Fig. S4A**) (Nik-Zainal et al., 2016).

**Fig. 4.**
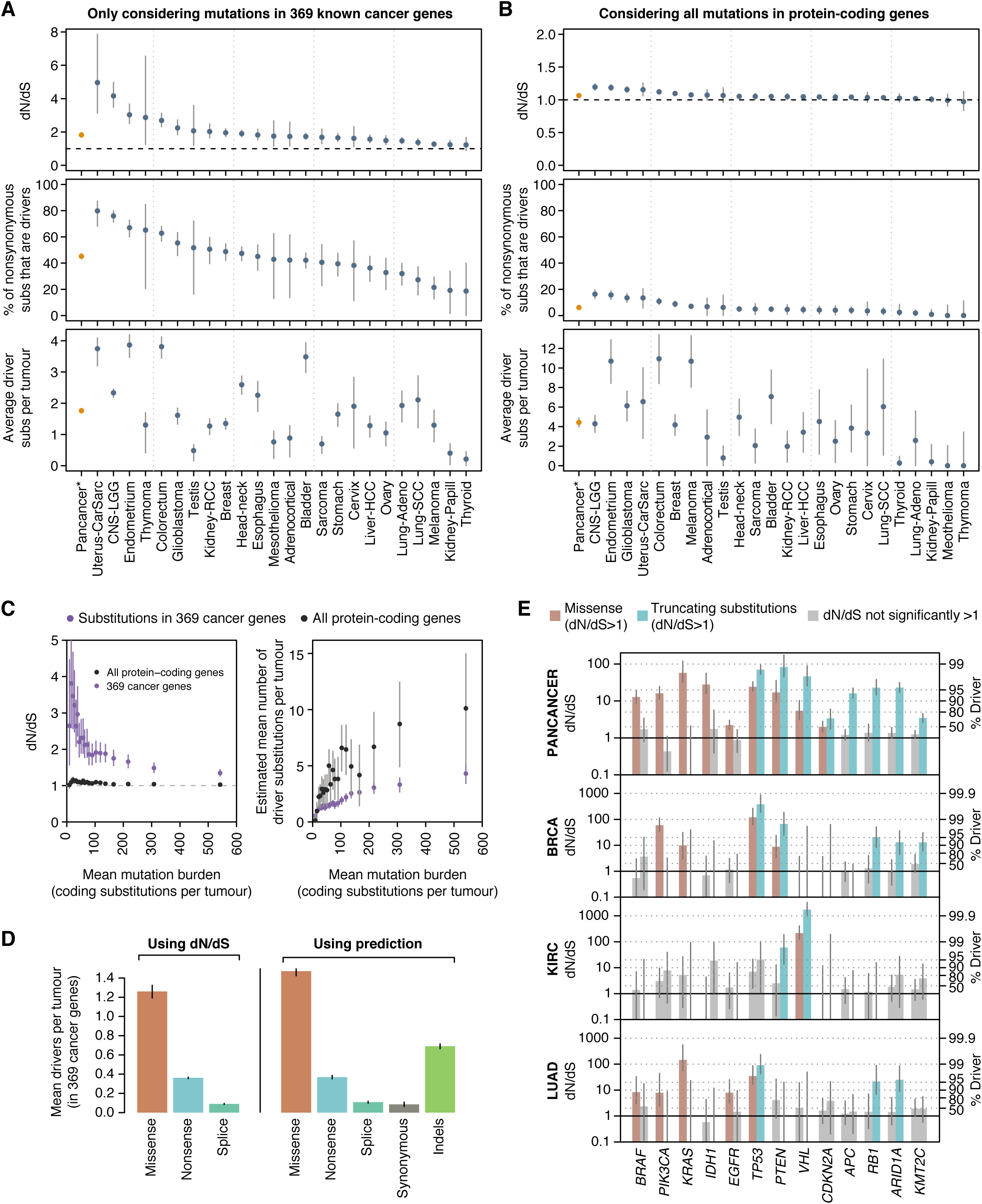
Estimated average number of driver mutations per tumor. (**A**) Top: global dN/dS values obtained for 369 known cancer genes. This analysis uses a single dN/dS ratio for all non-synonymous substitutions (missense, nonsense and essential splice site). Middle: percentage of non-synonymous mutations that are drivers assuming negligible negative selection. Bottom: average number of driver coding substitutions per tumor. Pancancer refers to the 24 cancer types with in-house mutation calls (Supplementary Methods S5). (**B**) Same panels as Fig 4A but including all genes in the genome. (**C**) dN/dS and estimated number of driver mutations per tumor grouping samples in 20 equal-sized bins according to mutation burden. This analysis excludes melanoma samples to avoid mutational biases (Supplementary Methods S5). Figures 4A-C were generated under the pentanucleotide substitution model for maximum accuracy. (**D**) Average number of driver mutations per tumor in 369 known cancer genes, using two different approaches: (1) dN/dS, (2) fitting a Poisson regression model with covariates on putative passenger genes and using this to measure the excess of mutations in known cancer genes (Supplementary Methods S5.3). This allows estimating the driver contribution of indels and synonymous mutations. (**E**) Left y-axis: dN/dS values for missense and truncating substitutions for a series of driver genes and for different datasets. Right y-axis: corresponding estimates of the fraction of driver mutations. Grey bars depict dN/dS ratios not significantly different from one. Error bars depict 95% confidence intervals.

Estimation of the number of driver mutations per tumor using this approach requires an accurate calculation of dN/dS ratios and so we took additional cautionary steps. Small inaccuracies in the mutation model can lead to systematic biases in the estimated numbers of drivers, especially in patients with high mutation burden. We found that this was particularly problematic in melanoma where the mutation signature is known to have sequence context biases beyond the immediate 5’ and 3’ neighbors of the mutated base (Pleasance et al., 2010a). A mutation model based on the pentanucleotide sequence context considerably outperformed the trinucleotide model in melanoma (Supplementary Methods S5.2.1 and Text S7). Reassuringly, for all other tumor types, estimates of the number of driver substitutions per tumor obtained under the trinucleotide and pentanucleotide models were highly consistent (**Fig. S1E**), indicating that possible unaccounted substitution biases are unlikely to impact our results.

Estimated dN/dS values on 369 known cancer genes varied extensively across cancer types (**Fig. 4A**). Using these ratios, we can estimate that 80% of the non-synonymous mutations occurring in cancer genes in low-grade glioma are driver mutations, whereas only 20% are drivers in melanoma, with other tumor types spanning this range. Combining these estimates with total mutation burden, we infer that the average number of coding substitutions in known cancer genes that are driver mutations ranges from <1/patient in sarcomas, thyroid cancers and mesotheliomas to 3-4/patient in bladder, endometrial and colorectal cancers (**Fig. 4A**).

Importantly, we can extend this analysis to all genes in the genome to provide the first comprehensive estimates of the total number of driver coding substitutions per tumor. Unlike simply counting the number of non-synonymous mutations seen in known cancer genes, this estimate is not constrained to known cancer genes and comprehensively measures the number of all coding driver substitutions per tumor. We find that the fraction of all coding mutations estimated to be drivers is low in most cancer types (**Fig. 4B**). For example, only 5.0% (CI_95%_: 3.0%-6.9%) of non-synonymous coding point mutations in head and neck cancers are predicted to be drivers. Interestingly, the average number of coding substitutions per tumor that are driver mutations is consistently modest, typically around 4/tumor and ranging from 1-10/tumor across tumor types (**Fig. 4B**). We note that this is an estimate of the *average* number of coding driver substitutions per tumor for each tumor type; the actual number for individual patients might vary extensively around this average. Estimates of the number of driver mutations per tumor based on all genes are about twice those from the 369 cancer genes, suggesting that about half of driver mutations occur in cancer genes yet to be discovered (**Fig. 4B**; **Fig. S4B**).

Mutator phenotypes are common in cancers, and it has been unclear whether their effect is to increase the overall number of driver mutations or simply to allow a clone to acquire a fixed complement of drivers faster than competing clones. We estimated the number of driver mutations per tumor as a function of total mutation burden, finding that as mutation burden increases, the dN/dS ratio converges towards 1 (**Fig. 4C**). When this is converted to estimate the number of coding substitutions that are drivers, we find that there is a hyperbolic relationship – as mutation burden increases, so too does the number of drivers, but not linearly. This implies that mutator phenotypes do indeed increase the overall number of driver mutations per tumor, even though they represent an ever smaller proportion of the total mutation burden.

The preceding estimates are limited to coding non-synonymous base substitutions. To extend this reasoning to estimate the numbers of small indels and synonymous substitutions that could be drivers, we measured the overall excess of these changes in known cancer genes by using putative passenger genes to estimate background mutation rates (Supplementary Methods S5.3) (Supek et al., 2014). Although these values are likely to be slight underestimates due to the small number of driver mutations hidden in undiscovered cancer genes, this will have minimal quantitative impact. Reassuringly, this more extensive model yielded very similar estimates for non-synonymous coding substitutions to these obtained above from dN/dS (**Fig. 4D**). We find that indels appear to contribute a similar number of driver mutations as truncating substitutions (nonsense and essential splice site mutations), with an average of ∼0.7 coding indel drivers per tumor in the 369 known cancer genes. Furthermore, synonymous driver mutations are rare but not negligible (∼0.09 per tumor in known cancer genes), in agreement with previous studies (Supek et al., 2014). For a more detailed analysis of the driver role of some synonymous mutations, see Supplementary Methods S5.3.1.

### Gene-by-gene, histology-by-histology driver mutations

Ultimately, if we are to use genomics to underpin precision medicine, an important step will be to infer which mutations in a given patient are drivers. As we have seen, not all somatic mutations in a given cancer gene are drivers, but dN/dS offers a framework to estimate these probabilities. We find that across tumor suppressor genes, whether missense substitutions are likely to be drivers or not varies considerably. For example, the tumor suppressors *ARID1A*, *RB1* and *APC* show dN/dS values for missense mutations close to one suggesting that the vast majority of missense mutations seen in these genes across all cancers are genuinely passengers, even though >95% of observed truncating mutations are estimated to be drivers (**Fig. 4E**). In contrast, the dN/dS value for missense mutations in *TP53* indicates that >95% of the missense mutations observed in this gene are drivers.

Such analyses highlight important differences across tumor types in the distribution of driver mutations. For example, in breast cancer, virtually all nonsense substitutions and ∼90% of missense substitutions in *PTEN* are driver mutations. However, in clear cell kidney cancer only nonsense mutations in *PTEN* are significantly enriched, with no significant excess of missense substitutions above expectation; and in lung adenocarcinoma, neither missense nor nonsense substitutions in *PTEN* were in statistical excess. Similarly, for oncogenes, we estimate that >10% of missense substitutions in *PIK3CA* in lung adenocarcinomas are passenger mutations, whereas only 1-2% of such events in breast cancer are (**Fig. 4E**).

## DISCUSSION

By adapting methods from evolutionary genomics and applying them to thousands of cancer genomes and to five healthy tissues, we have observed a universal pattern of selection in somatic evolution, characterized by a dominance of positive over negative selection. We have found that negative selection is a surprisingly weak force during cancer development, which in turn has allowed us to obtain genetic estimates of the number of driver coding substitutions per tumor.

Here, we have provided first exome-wide genetic estimates of the number of coding driver mutations across a range of tumor types. We have found that many tumors have 6 or more coding driver mutations, with endometrial cancer, colorectal cancer and melanoma averaging more than 10 per tumor. These counts must represent a lower bound for the true number of driver mutations, since they do not include non-coding driver point mutations nor structural variant drivers. Yet, overall the numbers of driver coding substitutions estimated per tumor are moderately low, more in line with classic models of cancer evolution (Armitage and Doll, 1954) than with recent speculations (Castro-Giner et al., 2015).

The absence of negative selection on coding point mutations in cancer is remarkable, especially since it is the predominant evolutionary pressure in the germline. Clearly, the vast majority of genes are dispensable for any given somatic lineage, presumably reflecting the buffering effect of diploidy and the inherent resilience and redundancy built into most cellular pathways. This explains why cancers can tolerate extreme levels of hypermutation, evidenced by tumors that acquire many hundreds of mutations with every cell division (Shlien et al., 2015). Our results also suggest that negative selection on point mutations is largely absent during normal somatic tissue maintenance as well. This has important implications for the somatic mutation theory of ageing (Morley, 1995), since it would argue that point mutations deleterious to the carrying cell do not drive cellular senescence, exhaustion and death. Rather, if point mutations do play a role in ageing of somatic tissues, it will be through the functional consequences to the organism of mutations that are selectively neutral or advantageous to the clone.

The conceptual framework we have developed for directly enumerating the excess or deficit of mutations over neutral expectation could be adapted to explore the role of driver mutations in non-coding regions of the genome. Furthermore, with increasing numbers of tumors being sequenced, we will be able to deploy such reasoning at ever higher resolution to estimate probabilities that variants in particular exons or domains of a gene in a particular tumor type are driver mutations. Such approaches could ultimately underpin statistically rigorous, personalized annotation of driver mutations, a crucial step in successfully implementing precision oncology.

## Author contributions

IM developed the statistical methodology, analyzed and interpreted the data. IM, KJD, KMR, KH and PVL downloaded the TCGA data and called somatic mutations. HD annotated driver mutations in breast cancer. MG, MRS and PJC provided key advice. PJC supervised the project. IM and PJC wrote the manuscript.

## Acknowledgements

IM is supported by EMBO and Cancer Research UK. P.V.L. is a Winton Group Leader in recognition of the Winton Charitable Foundation’s support towards the establishment of The Francis Crick Institute. PJC is a Wellcome Trust Senior Clinical Fellow. This work was also supported by the Wellcome Trust and the Francis Crick Institute, which receives its core funding from Cancer Research UK (FC001202), the UK Medical Research Council (FC001202), and the Wellcome Trust (FC001202). We thank Matt Hurles for comments and suggestions. The results published here are based upon data generated by the TCGA Research Network:

*http://cancergenome.nih.gov/*.

An R package with the code to run *dNdScv* is in preparation and will be made publicly available with this publication.

## Supplementary Materials for Universal patterns of selection in cancer and somatic tissues

### 1. A dN/dS model for cancer genomics

dN/dS (also called Ka/Ks) is the ratio between the rate of non-synonymous substitutions per non-synonymous site and the rate of synonymous substitutions per synonymous site. First developed in the 1980s (Miyata and Yasunaga, 1980; Nei and Gojobori, 1986), it has a long history in the detection of negative and positive selection from sequencing data (Yang and Bielawski, 2000).

dN/dS is particularly suitable for the analysis of coding mutations in cancer genomes, for several reasons. First, unlike evolutionary comparisons of distant species, in which a change between two sequences may be the result of multiple changes to the site over the course of evolution, the density of substitutions per site in cancer is extremely low (typically <10^-5^ mutations per site) (Martincorena and Campbell, 2015). This greatly simplifies the estimation of rate parameters and facilitates the development of more complex mutation and selection models (Greenman et al., 2006). Second, while some concerns exist regarding the use of dN/dS within a highly-recombining population (Kryazhimskiy and Plotkin, 2008), these considerations do not apply to somatic mutations accumulated in a cancer sample. That is both because cancer cells evolve asexually and because collections of somatic mutations are identified by comparing a cancer sample to the ancestral genome, rather than comparing two individuals or cells from a population. Finally, dN/dS offers a measure of selection largely free of assumptions, in contrast to population genetic tests of selection, in which apparent violation of neutrality can result from demographic changes rather than selection.

#### 1.1 Poisson framework

In this study we adopt and expand upon the Poisson framework developed by Greenman *et al*. (Greenman et al., 2006). Mutations are classified according to their substitution type (*i*) (depending on the substitution model) and functional impact (synonymous –*s*-, missense –*m*-, nonsense –*n*-and essential splice sites –*e*-). Note that, throughout the paper, the term “truncating substitutions” refers to nonsense and essential splice site substitutions together. For example, the number of C>T synonymous mutations (*n*_C>T,s_) in a collection of samples is modelled as a Poisson process:

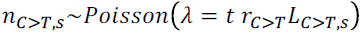

Where *t* is the density of substitutions per site, *r*_C>T_ is the relative rate of C>T substitutions per site, and *L*_C>T,s_ is the number of C sites in which a C>T change is synonymous. In this parameterization, one rate parameter of the substitution matrix is arbitrarily set to 1 (*e.g*. r_G>T_=1, so that all other rates are relative rates with respect to it). For non-synonymous sites, an extra parameter reflects the effect of selection on the accumulation of mutations:

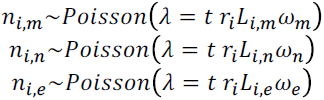

The ω parameters are the dN/dS ratios inferred by the model after correcting for the rates of different substitution classes (*r*_i_) and for sequence composition (*L*). Maximum-likelihood estimates for all parameters in the model can be efficiently obtained by Poisson regression.

Although a Poisson implementation of dN/dS is particularly suitable for cancer genomic data, it can similarly be used in other resequencing studies, especially as long as the density of mutations per site is low. This includes, for example, studies of human evolution and bacterial populations.

#### 1.2 Substitution models

The simplest dN/dS implementations, such as the Nei-Gojobori model (Nei and Gojobori, 1986), treat all substitutions as a single substitution class. More sophisticated likelihood-implementations, widely used nowadays, instead use a substitution model with two substitution classes: transitions (C<>T, A<>G) and transversions (C<>A, C<>G, T<>A, T<>G) (*i.e.* they use a transition/transversion ratio as a single rate parameter) (Goldman and Yang, 1994). More complex mutation models include the GTR (General Time Reversible) model with 6 mutation classes, one for each of the 6 possible reversible base changes.

Somatic mutations in cancer have been shown to display strong context-dependence, particularly from one base upstream and downstream of the mutant base (Alexandrov et al., 2013). As we show in Supplementary Text S6 and Fig. S1A, the use of simplistic substitution models can lead to severe systematic under- or over-estimation of dN/dS ratios and erroneous inference of selection. Previous studies of selection in cancer genomics have accounted for only some of this context-dependence, especially the high rate of C>T at CpG dinucleotides (Greenman et al., 2006; Lawrence et al., 2013; Yang et al., 2003).

In this study, to comprehensively avoid biases emerging from context-dependent effects from one base upstream and downstream of the mutant base, we use a full trinucleotide model with 192 rate parameters, one for each of the possible trinucleotide substitution rates. By using a model with 192 rates, as opposed to 96 rates, we accommodate the possibility of strand asymmetry emerging from transcription coupled repair in coding regions (Pleasance et al., 2010b). More complex models, including a full pentanucleotide substitution model, were also evaluated for specific applications (see Supplementary Methods S5.2.1, Supplementary Text S7 and Fig. S1D,E).

#### 1.3 Modeling variable mutation rates across genes: *dNdScv*

In early exome studies with small numbers of samples, methods to detect significant mutation recurrence at gene level often assumed that the substitution rate was uniform across genes (Bolli et al., 2014; Greenman et al., 2007; Lawrence et al., 2013). In the Poisson framework described above, this is achieved by having a single *t* parameter shared across all genes (***uniform rate dN/dS model***). Maximum-likelihood estimates for the parameters across genes (*t*, *r*_i_, ω_m_, ω_n_ and ω_e_) are obtained by Poisson regression.

However, mutation rates are known to vary substantially across genes and models assuming uniform mutation rates across genes lead to the identification of large numbers of false positives when applied to relatively large numbers of samples (Lawrence et al., 2013). A simple way to avoid this problem is to have a separate *t* parameter for each gene (***variable rate dN/dS model***). This is similar to most dN/dS implementations used in comparative genomics, in which the background mutation rate in a gene is directly estimated from the number of synonymous mutations observed in the gene. Although we have used this model successfully in cancer genomic datasets containing thousands of samples (Wong et al., 2014), it lacks statistical power to detect positively selected genes in smaller datasets.

The mutation rate is known to vary across genes depending on their expression level, replication time and chromatin state (Lawrence et al., 2013; Pleasance et al., 2010a; Polak et al., 2015; Schuster-Bockler and Lehner, 2012). Some methods designed to identify recurrently mutated genes in cancer genomes, exploit this knowledge to improve their background mutation rate models. For example, *MutSigCV* uses three covariates to estimate the mutation rate of each gene, by using information from other genes with similar covariate values (Lawrence et al., 2014; Lawrence et al., 2013). Inspired by this work, we developed *dNdScv*, a method that combines dN/dS with a negative binomial regression on a large number of covariates.

We model the variation of the normalized mutation rate per base pair (*t*) across genes as following a Gamma distribution. In a given dataset, the observed number of synonymous mutations per gene –*j*-(*n*_s,j_) can then be modelled as a Poisson process whose mean is drawn from a Gamma distribution reflecting the variation of the mutation rate across genes.

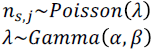

Since the negative binomial distribution is a Gamma-Poisson compound distribution, the number of synonymous mutations per gene is modelled as following a negative binomial distribution. This enables the use of a negative binomial regression framework to estimate the background mutation model across genes for each dataset. Gene size, gene sequence and the impact of the substitution model are all accounted for as an offset in the model (reflecting the *exposure* of the gene). The normalized mutation rate per site, *t*, is modelled as Gamma-distributed across genes, reflecting the uncertainty in the variation of the mutation rate across genes remaining after accounting for the exposure of the gene. Covariates can then be used in this framework, to improve the estimated background rate for a gene and reduce the unexplained variation of the mutation rate, and so reduce the dispersion of the underlying Gamma distribution. A reduction in the unexplained variation of the mutation rate leads to more sensitivity for the detection of selection, while the use of overdispersion in the form of the Gamma distribution, reflecting the uncertainty in mutation rates across genes, ensures good specificity.

In R code, the regression is performed using:

*model = glm.nb(n_syn ∼ offset(log(expected_syn)) + covariate_matrix)*

where:

*n_syn* for gene *j* is: *n*_s,j_ = ∑_i_ n_i,s,j_

*expected_syn* for gene *j* is: *E*_s,j_ *=∑_i_r_i_L_i,s_*(with ***t*** being constant across genes)

This framework allows to use a large number of covariates and variable selection approaches to improve the background mutation rate model. In this study, we have used as the covariate matrix the first 20 principal components of 169 chromatin marks from the RoadMap Epigenomics Project (Kundaje et al., 2015). This included data from 63 cell lines and 10 different epigenetic marks (H3K9me3, H3K36me3, H3K27me3, H3K4me1, H3K4me3, H3K9ac, H3K23ac, H3K14ac, H2AK9ac and DNase). Since it has been shown that epigenomic landscapes derived from cell lines more closely related to a cancer type are better predictors of its local mutation density (Polak et al., 2015), there is added value in using a wide set of epigenomic covariates. The use of a regression framework hence allows to build complex and fully data-driven background mutation models for each dataset.

The negative binomial regression estimates a Gamma distribution for the uncertainty on *t*j after considering the gene size, the gene sequence, the substitution model and the covariates. Hence, the likelihood for *t*_j_ can now be constrained both by the global knowledge of how the mutation rate varies across genes and the local number of synonymous mutations in the gene.

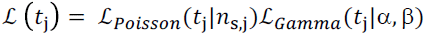

By using this joint likelihood, *dNdScv* weighs the amount of information on the mutation rate of the gene. In small datasets, in which most genes have zero or a few synonymous mutations, the Gamma function dominates the likelihood. In large datasets with sufficient numbers of synonymous mutations per gene, the Poisson function dominates and the model converges to the *variable rate dN/dS model*.

Derivation of the expression for 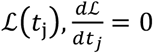, gives a simple analytical solution for the maximum likelihood estimate of *t*j under both the Poisson and Gamma constraints. The maximum likelihood estimate for the expected number of synonymous mutations in a gene under the *dNdScv* model 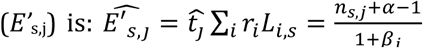. Where α and *β*_j_ are the shape and rate (inverse of scale) parameters of the Gamma distribution respectively, defined as: α=θ and *β*_j_=θ*/*µ_j_ (µ_j_ is the predicted number of synonymous mutations for gene *j* according to the negative binomial regression model and θ is the overdispersion parameter of the regression model).

Confidence intervals for ω parameters under the *dNdScv* model (as used in Fig. 4E) were obtained by profile likelihood integrating out *t*_j_.

#### 1.4 Likelihood ratio tests for the inference of selection

In all three dN/dS models (*uniform rate*, *variable rate* and *dNdScv*), inference of selection is performed using Likelihood Ratio Tests, similarly to traditional likelihood dN/dS models used in phylogenetics (Goldman and Yang, 1994; Yang and Bielawski, 2000). Examples of null and alternative hypotheses for different tests are shown below.

Global test for selection with free ω parameters (3 degrees of freedom):

H_0_: ω_m_ = 1; ω_n_ = 1; ω_e_ = 1

H_1_: ω_m_ ≠ 1; ω_n_ ≠ 1; ω_e_ ≠ 1

Global test for selection with a single ω parameter for truncating substitutions (nonsense and essential splice site mutations) (2 degrees of freedom). This is the test used in the screen for positively selected genes in this study as it tends to be more sensitive than the fully unconstrained model above.

H_0_: ω_m_ = 1; ω_n_ = 1; ω_e_ = 1

H_1_: ω_m_ ≠ 1; ω_n_ = ω_e_ ≠ 1

Test for selection on missense mutations (1 degree of freedom).

H_0_: ω_m_ = 1; ω_n_ ≠ 1; ω_e_ ≠ 1

H_1_: ω_m_ ≠ 1; ω_n_ ≠ 1; ω_e_ ≠ 1

Multiple testing correction is performed using Benjamini and Hochberg’s false discovery rate (Benjamini and Hochberg, 1995) for all genes tested. To boost the statistical power to detect selection on known cancer genes, we use *restricted hypothesis testing* on an *a priori* list of known cancer genes, as described before (Lawrence et al., 2014). In this study, we use the list of 174 *COSMIC classic genes* from version 73 of the COSMIC database (Forbes et al., 2015) for RHT in the positive selection screen, and a list of essential genes for RHT in the negative selection screen (see Supplementary Methods S4.4). Alternative approaches that can help increase power on *a priori* gene candidates without theoretically incurring in an inflation of the global false discovery rate are *stratified false discovery rate* (Sun et al., 2006) and *data-driven hypothesis weighting* (Ignatiadis et al., 2016).

#### 1.5. Recurrence of insertions and deletions

dN/dS can be used to detect and quantify selection on coding substitutions, but not on small insertions or deletions (indels). To identify genes recurrently affected by indels or by other mutation types, such as dinucleotide substitutions or complex substitutions, we use a different model.

Briefly, a simple negative binomial regression model is used to estimate the expected rate of indels per gene. The length of the CDS of each gene is used as an offset and the 20 epigenomic covariates used in *dNdScv* are also used as covariates here (Supplementary Methods S1.3). To minimise the risk of driver indels inflating the background model, known cancer genes are excluded when fitting the negative binomial model (in this study we used the list of 558 cancer genes in the *Cancer Gene Census* version 73 (Forbes et al., 2015). Applying this regression model to all genes in the genome provides an estimate of the mean indel rate expected in each gene and of the overdispersion of the model (θ). A *P*-value for the observed number of indels in each gene (*n*i,j) can be obtained using the cumulative negative binomial distribution. For each gene, we used Fisher’s method to combine the *P*-value from the indel model with the *P*-value obtained from *dNdScv* (with 2 degrees of freedom) for selection on coding substitutions. The resulting global *P*-value was used to identify genes under positive selection in Fig. 2 and Supplementary Table S2.

Supplementary Text S9 compares the specificity and sensitivity of the three dN/dS models introduced here (uniform rate dN/dS model, variable rate dN/dS model and *dNdScv*).

### 2. Exome data and mutation calling

#### 2.1. Exome data

Paired tumor and normal exome sequencing files from 9,699 cancer patients were downloaded from CGHub between November and December 2015. The samples, sequenced by The Cancer Genome Atlas (*http://cancergenome.nih.gov/*), correspond to 29 different tumor types. Colon and rectal cancer were grouped together as colorectal cancer. Supplementary Table S1 shows the list of cancer types used in this study, their TCGA 4-letter code names, the longer abbreviations used in this study and the number of samples eventually selected for analysis (7,664 across all cancer types).

#### 2.2. Calling of point mutations and indels

The data were uniformly reprocessed using the Wellcome Trust Sanger Institute’s variant calling algorithms to ensure uniformity across cancer types and to have control over the filtering of mutations at polymorphic sites. Owing to negative selection during human evolution, germline polymorphisms are heavily enriched in synonymous substitutions (Fig. 1A). As a consequence, incomplete removal of germline polymorphisms from the collections of somatic mutations can lead to an underestimation of dN/dS ratios, while removal of genuine somatic mutations at polymorphic sites can lead to an overestimation of dN/dS ratios (see Supplementary Text S8 and Fig. S1B,C for analyses on the impact of germline SNPs in catalogs of somatic mutations).

Paired-end reads were aligned to the reference human genome (GRCh37, hs37d5 build) using *BWA-MEM* (Li, 2013). Substitutions were called using *CaVEMan* (Cancer Variants Through Expectation Maximization: http://cancerit.github.io/CaVEMan/) (Jones et al., 2016). Indels were called using cgpPindel v2.0 (http://cancerit.github.io/cgpPindel/) (Raine et al., 2015). A panel of unmatched normal samples (sequenced at the Wellcome Trust Sanger Institute) was used to remove common sequencing and mapping artefacts.

#### 2.3 Quality controls and use of TCGA calls in five cancer types

Only pairs of samples with the same TCGA barcode ID to those used by TCGA in their public somatic mutation calls were considered for further study. To minimize the risk of germline polymorphisms in the collections of somatic mutations, somatic calls at sites with less than 10 reads of sequencing coverage in the matched normal sample were excluded. To ensure that somatic calls from our pipeline were not excessively different from those released by TCGA, samples in which our algorithms called <50% of the coding mutations publicly released by TCGA were excluded. Samples with >3,000 coding mutations (*i.e.* ∼100 mutations/Mb), including substitutions and indels, were excluded from all of the analyses in this study. After applying these filters, a total of 7,664 samples were used for the analyses in this paper.

Comparison of the mutation calls obtained from our pipeline to those released by TCGA for the same samples suggested low sensitivity of our pipeline in five of the 29 cancer types analyzed: acute myeloid leukemia (LAML), kidney chromophobe (KICH), pheochromocytoma and paraganglioma (PCPG), prostate adenocarcinoma (PRAD) and pancreatic adenocarcinoma (PAAD). For these five cancer types, public TCGA mutation calls were used in this study instead of those from our pipeline. These five cancer types were used in the driver discovery analyses, since these analyses are largely robust to minor germline contamination or over-filtering at polymorphic sites. However, these cancer types were excluded from the analyses of negative selection (Fig. 3, Supplementary Methods S4) and the estimation of number of driver substitutions per tumor (Fig. 4, Supplementary Methods S5), where moderate biases to dN/dS can affect the interpretation of the results.

#### 2.4. Calling of copy number changes

We used the ASCAT algorithm (Van Loo et al., 2010) to identify copy number changes across 13,241 TCGA samples using Affymetrix SNP6 arrays. CEL files provided by TCGA were processed using PennCNV libraries (Wang et al., 2007) to obtain logR and BAF values. The logR values were subsequently corrected for GC content to decrease wave artefacts, which often affect samples profiled by SNP arrays. Copy number profiles for all tumor samples were then inferred from the corrected data using ASCAT version 2.4.2 (Van Loo et al., 2010).

### 3. Screen for positive selection at gene level (driver gene discovery)

To identify genes under significant positive selection we ran *dNdScv* on every cancer type separately and on all 7,664 samples together. *P*-values were calculated as described in Supplementary Methods S1.3-1.5 and adjusted for multiple testing using Benjamini and Hochberg’s false discovery rate (Benjamini and Hochberg, 1995). On inspection of the results, a small number of significant genes were found to be false positives resulting from recurrent sequencing or mapping artefacts in the collections of somatic mutations. To systematically remove false positives due to recurrent artefacts, all mutations found in significant genes were subject to an *in silico* validation (see below), false calls were removed and *dNdScv* rerun on the cleaned dataset.

Genes found as significant (q-value<0.05) in each cancer type are depicted in Fig. 2 and in Supplementary Table S2. Since combining results from multiple tumor types can inflate the global false discovery rate in the final list of significant genes, we then performed a global multiple testing correction on the entire matrix of *P*-values (20090 genes by 30 datasets) (as in (Lawrence et al., 2014)). This resulted in a list of 180 putatively positively-selected (driver) genes. Using restricted hypothesis testing led to the additional identification of 24 driver genes (Supplementary Methods S1.4) (Lawrence et al., 2014).

#### 3.1. *In silico* identification and removal of sequencing artefacts

Evaluation of significant hits revealed a small number of false positives due to recurrent artefacts that escaped our filters and our unmatched normal panel. To systematically identify recurrent artefacts leading to false positives in the screens for positive and negative selection, we used *ShearwaterML* (Gerstung et al., 2014; Martincorena et al., 2015).

*ShearwaterML* is a variant calling algorithm that relies on building a base-specific error model by using a large collection of unmatched normal samples. Sequencing artefacts caused by *Illumina* sequencing errors, PCR errors, DNA damage in a library, misalignment of reads or other causes, are expected to appear at similar frequencies in sequencing libraries of tumor or healthy (normal) tissue. Thus, all mutations identified in genes detected as significant by *dNdScv* were re-evaluated by *ShearwaterML*, comparing the number of reads supporting the mutation in the mutant sample to the frequency of errors seen across a large panel of TCGA normal samples from the same cancer type using a beta-binomial likelihood model (Martincorena et al., 2015).

To build a reliable panel of normal samples for each TCGA dataset and avoid filtering out genuine driver mutations, we excluded from the panels any normal sample with suggestive evidence of a mutation (>=3 supporting reads) in a list of 344 recurrently mutated sites in known cancer genes. This reduces the risk of including samples in the normal panel with significant tumor contamination or hematopoietic clonal expansions (Xie et al., 2014).

*P*-values resulting from *ShearwaterML* were adjusted for multiple testing using Benjamini and Hochberg’s false discovery rate (Benjamini and Hochberg, 1995), correcting for n=*N*\**S* tests to avoid a discovery bias (where *N* is the number of sites tested and *S* is the number of samples in each cancer type). Mutations with q-value>0.20 were removed and *dNdScv* was re-run on the cleaned dataset. 49 genes were found to be heavily affected by artefacts, with more than 50% of the mutations found in them being considered artefactual by *ShearwaterML*. These genes were conservatively excluded from any significant hits in the positive selection screen.

The 49 genes heavily-affected by artefacts are: *AGAP10, AL445989.1, ANAPC1, ANKRD36C, AQP7, BMI1, C16orf3, CD209, CDC27, CDC7, CRIPAK, DTD2, EP400, FAM104B, FRG1, FRG1B, GNAQ, HLA-DRB5, HSPD1, IGBP1, KBTBD6, KRT14, KRT5, KRT6A, KRTAP1-5, KRTAP4-11, KRTAP4-3, KRTAP4-8, KRTAP4-9, KRTAP5-5, KRTAP9-9, MLLT3, MUC4, MUC8, NCOA6, PABPC1, PCDHB12, POTEC, POTEM, PPFIBP1, PRKRIR, PTH2, RGPD3, RGPD8, RP11-176H8.1, SLC35G6, TMEM219, TPT1* and *UBBP4.*

### 4. Negative selection analyses

#### 4.1 Samples selected for negative selection analyses

As we have described, simplistic mutation models, germline contamination of the catalogs of somatic mutations and over-filtering of genuine somatic mutations at polymorphic sites can lead to biased dN/dS ratios. When analyzing dN/dS ratios close to 1, these biases can lead to wrong inferences about selection, as shown in Fig. S1A-E and Supplementary Text S6-8.

In order to avoid these biases, the analyses of negative selection shown in Fig. 3 were carried out on a subset of all samples, encompassing 5,763 samples from 23 cancer types. First, the five cancer types with TCGA mutation calls were excluded from the analyses to have control over the filtering of germline mutations used during variant calling. Second, melanoma samples were excluded from these analyses since the mutation spectrum in melanoma causes a downward bias to dN/dS under the trinucleotide model (Supplementary Text S7). Third, only samples with copy number information were included in the analyses, since this information was required for several of the analyses. Finally, only samples with fewer than 500 coding mutations per exome were included in the analyses to avoid hypermutator samples dominating the analyses and ensure representative results.

#### 4.2. dN/dS distributions across genes

Observed dN/dS values at gene level are subject to considerable uncertainty due to the limited number of substitutions per gene. Hence, the variation in dN/dS values observed across genes (Fig. 3A) is a composite of the true variation of selection across genes and Poisson noise in the counts of non-synonymous and synonymous mutations. Using mixture models, this technical variation can be eliminated to infer the underlying dN/dS distribution across genes.

For any gene, given a extent of selection (ω_m,j_) and a substitution model, the expected fraction of synonymous and missense substitutions in the gene can be calculated as follows: 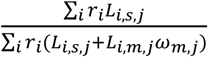, 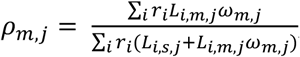 respectively. The analysis of truncating substitutions (nonsense and essential splice site mutations) was done analogously.

##### 4.2.1. Neutral simulations

To study how much variation in observed dN/dS values across genes is expected by simple noise under perfect neutral evolution, we first carried out a simple simulation. Using the expected fraction of synonymous and missense mutations per gene under neutrality (ρ_s,j,neutral_ and ρ_m,j,neutral_ given ω_m,j_=1 for all genes), and the total number of mutations observed per gene (*n*s,j+*n*m,j), we performed a random binomial simulation of the number of missense mutations per gene: *n*_*m,random*_∼*B*(*n = n*_*s,j*_ + *n*_*m,j*_,*p* = ρ_*m,j,neutral*_) This yields a maximum-likelihood point estimate for dN/dS per gene of:(*n*_*m,j,random*_.ρ_*s,j,neutral*_)/(*n*_*s,j,random*_.ρ_*m,j,neutral*_) This simulation revealed that most of the apparent variation observed in dN/dS across genes was technical, caused by the limited number of mutations per gene (Fig. 3A).

##### 4.2.2. Binomial mixture model

We can go beyond neutral simulations and infer the extent of the biological variation of ω across genes. Given *ρ*_s,j_ and *ρ*_m,j_ (as a function of ω_m,j_) and the total number of mutations seen in the gene (*n*_s,j_+*n*_m,j_), the probability of observing *n*_m,j_ mutations in a gene follows a binomial distribution. The advantage of using a binomial distribution contingent on the total number of observed mutations is that it makes the approach unaffected by the uncertainty in the background mutation rate of the gene.

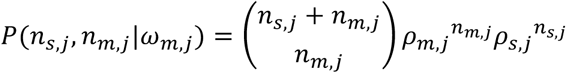

We can extend this to model ω as a distribution across genes, for example using a discrete mixture model or integrating over a continuous distribution for ω. This is similar to existing approaches for modeling the distribution of ω across codons of a protein in comparative genomics (Nielsen and Yang, 1998). In this study, to avoid imposing a restrictive parameterization of the distribution of ω (dN/dS) across genes, we used a flexible discrete distribution with a fine grid. For the results shown in the main text, we used a discrete distribution defined as ω ∈ (0, 0.1, 0.2, 0.3, 0.4, 0.5, 0.6, 0.7, 0.8, 0.9, 1, 1.1, 1.2, 1.3, 1.4, 1.5, 1.6, 1.7, 1.8, 1.9, 2, 3, 4, 5, 10, 15, 20), with a free probability mass function defined at these values 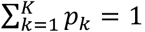, where K is the number of points used in the discrete distribution and *p*_k_ the fraction of genes with ω*=*ω*_k_*). The global likelihood of the distribution of ω across genes can then be expressed as:

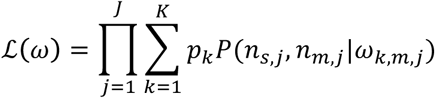

Where *J* is the total number of genes considered. Maximum likelihood estimates for the probability mass function 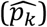 were obtained using an Expectation Maximization (EM) algorithm, initialized with uniform probabilities (*p*_k,0_ = 1/K). Confidence intervals were obtained by bootstrapping (sampling genes with replacement). Using distributions with more points yielded analogous results.

##### 4.2.3. Estimation of the average number of mutations under positive and negative selection per tumor

Once the underlying ω distribution has been estimated 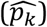, the probability that a gene has a particular value of ω can be calculated using the equation below. This equation corresponds to the posterior probability of a gene belonging to a ω class in the EM algorithm and it is identical to the empirical Bayes equation used for a similar purpose in the dN/dS literature (Nielsen and Yang, 1998).

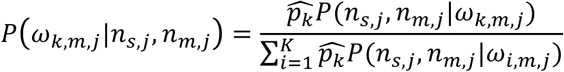

If a gene is evolving under a given value of ω_k,m,j_, the maximum likelihood estimate for the expected number of missense mutations in the gene, given the mutations observed, is: 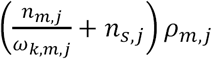 If we assume that genes under positive selection (ω_k,m,j_>1) do not contain a significant number of sites under negative selection, or vice versa, we can use the value of ω_k,m,j_ to estimate the number of missense mutations fixed by positive selection or depleted by negative selection. Summing over all genes, we can obtain global estimates for the average number of missense mutations fixed by positive selection (δ_pos_) or depleted by negative selection (δ_neg_), per tumor.

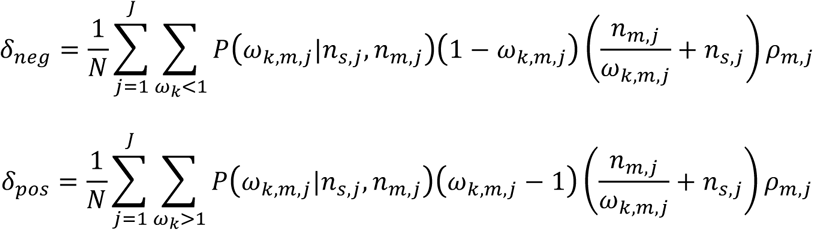

Where *N* is the number of samples used in the analysis. Confidence intervals for these estimates were obtained by bootstrapping (sampling genes with replacement). It should be noted that, in the presence of both positive and negative selection acting on the same gene at different sites or in different samples, these estimates will underestimate the extent of positive and negative selection. However, in order to explain the observation that the vast majority of genes are estimated to have an average ω∼1 (Fig. 3B), the combination of positive and negative selection should be nearly perfectly balanced across most genes in the genome, which is unlikely. This suggests that most genes seem to accumulate missense mutations largely neutrally and so that δ_pos_ and δ_neg_ are probably decent approximations.

In this study, we inferred 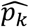, δ_pos_ and δ_neg_ for missense and truncating (nonsense and essential splice site) substitutions separately, as well as for three classes of mutations according to the copy number state of the region where the mutations occurred: haploid regions (1:0), loss of heterozygosity (LOH) regions with higher ploidy (n:0, with n>1), and all others (*i.e.* regions without LOH). Estimates shown in Fig. 3D include the sum of all of these mutation types. Fig. 3A-C show estimates for missense mutations in regions without LOH.

#### 4.3. Gene-level analyses of negative selection

To identify whether any gene has a dN/dS value significantly lower than 1 we used a one-sided test on missense mutations alone:

H_0_: ω_m_ = 1; ω_n_ ≠ 1; ω_e_ ≠ 1

H_1_: ω_m_ ≤ 1; ω_n_ ≠ 1; ω_e_ ≠ 1

This test has 1 degree of freedom and the resulting *P-*value from the Chi-square distribution is divided by two, as the test is one-sided. Since tissue-specific datasets lack statistical power to detect negative selection at gene level, we used the entire pancancer dataset (n=7,664 samples) for this analysis. We used both the *dNdScv* model and the *variable rate dN/dS model* (Supplementary Methods S1.3). These tests did not find any gene under significant negative selection at false discovery rate <10%.

To boost the statistical power on genes that may be suspected to be under stronger negative selection, we performed restricted hypothesis testing on an *a priori* chosen list of 1,734 essential genes (see Supplementary Methods S4.4 below for a description of the genes in this list). All genes yielded q-values higher than 0.10.

##### 4.3.1. Power calculations

The power to detect negative selection in a gene (or a group of genes) under the *variable rate dN/dS model* is determined by two main factors: (1) the effect size (the dN/dS ratio), and (2) the number of mutations in the gene (which is largely determined by the number of samples in the dataset, their mutation burden and the length of the gene). Under the *dNdScv* model, a third factor affecting the power is the uncertainty of the background model (*i.e.* the overdispersion of the negative binomial regression -θ-).

In order to study the power to detect negative selection under both models, we performed random simulations. Let ***m*** be the expected (average) number of coding mutations in a gene in a dataset, *ρ*_s_, *ρ*_m_ and *ρ*_t_ the fraction of synonymous, missense and truncating (nonsense and essential splice site) substitutions expected under neutrality, ω_m_ and ω the corresponding values of dN/dS, and α the shape parameter of the underlying Gamma distribution. Then we can simulate the number of synonymous, missense and truncating substitutions in the gene using:

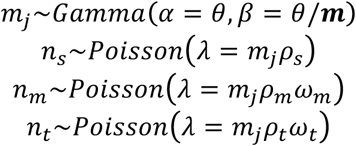

*P*-values for two-sided tests under both the *variable rate dN/dS model* and *dNdScv* can be calculated from these numbers as described in Supplementary Methods S4.3 (H_0_: ω_m_=1; H_1_: ω_m_≠1, df=1). For each combination of ***m*** and ω tested, we performed 5,000 simulations. The fraction of *P-*values below 0.05 reflect the power to detect a gene as significantly under selection. The values used for *ρ*_s_, *ρ*_m_, *ρ*_t_ and θ are the average values for these parameters observed in the pancancer dataset (*ρ*_s_=0.287, *ρ*_m_=0.649, *ρ*_t_=0.064, θ=6.03).

#### 4.4. Group-level analyses of negative selection

Given the limited statistical power to detect negative selection at the level of individual genes, we searched for evidence of negative selection in groups of related genes. To do so, we first excluded a long list of 987 putative cancer genes, by combining gene lists from multiple sources. We then used the *variable rate dN/dS model* to study selection on groups of genes, as defined by expression level, local chromatin state, essentiality and gene ontology functional annotation.

*Expression*: As a measure of the typical expression level of a gene across tumors, we calculated the mean of the log RSEM-normalized expression level of each gene across a collection of 6,190 TCGA samples (*.rsem.genes.normalized_results* TCGA files).

*Chromatin state*: As a measure of the typical chromatin state of a gene, we defined as heterochromatin and euchromatin those regions in which the six main ENCODE cell lines shared the same annotation (ENCODE Project Consortium, 2012).

*Essentiality*: As a list of genes essential for cell survival and growth, we used a collection of 1,734 core essential genes reported by a recent mutagenesis screen in haploid human cell lines (Blomen et al., 2015). This list of genes is heavily enriched in proteins participating in key cellular components and pathways, such as the ribosome, the spliceosome, the aminoacyl-tRNA biosynthesis pathway, the proteasome, RNA degradation, DNA replication, RNA polymerases and the cell cycle (Blomen et al., 2015).

*Gene ontology*: To search for evidence of negative selection at the level of functionally related genes, we used *Ensembl BioMart* to extract Gene Ontology (GO) annotations for all genes. To ensure adequate statistical power and reduce multiple testing correction, we only tested groups composed of at least 30 genes. We considered GO annotations of *Biological Processes*, *Cellular Components* and *Molecular Functions*. Overall we tested 1,242 functional groups of genes and performed Bonferroni multiple-testing correction (we used Bonferroni to account for the extensive overlaps between gene groups). Including all genes in the analysis yielded a large number of GO groups with evidence of positive selection on missense and/or nonsense substitutions (n=428), but no group with evidence of negative selection. Excluding the long list of 987 putative driver genes dramatically reduced the number of functional gene groups with evidence of positive selection (n=27), but still no GO group showed evidence of significant negative selection (Fig. S3). Repeating this analysis on mutations occurring in haploid regions did not identify any group of genes under clear negative selection.

### 5. Estimation of the number of driver mutations

#### 5.1. Samples selected for the estimation of the number of driver mutations

All samples with *CaVEMan* mutation calls and less than 500 coding mutations per sample were included in this analysis, including the melanoma dataset. Overall, a total of 6,108 samples from 24 cancer types were included in the pancancer estimates of the number of driver mutations per tumor shown in Fig. 4.

#### 5.2 Estimating the number of substitutions fixed by positive selection from dN/dS

In the absence of negative selection and mutation biases, we can accurately estimate the number of mutations expected to have accumulated neutrally in a gene or group of genes. As described by Greenman *et al*. (Greenman et al., 2006), this can be used to estimate the number of mutations in excess that have been fixed by positive selection. Assuming a negligible role for negative selection, we can calculate the fraction (*f*_m_) and the absolute number (*d*_m_) of mutations in a gene or group of genes that are genuine driver mutations as:

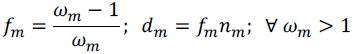

In the presence of significant negative selection, these equations would provide lower bound estimates of the density and number of genuine driver mutations. However, our analyses suggest that negative selection has a small quantitative effect on the accumulation of passenger mutations in cancer. The same equations apply for nonsense and essential splice site substitutions.

Naively, one might expect that all non-synonymous mutations observed in a driver gene could have been positively selected and so that they could all be drivers. However, in the absence of negative selection, we should still expect passenger mutations to accumulate in driver genes at approximately the background rate predicted under neutrally and so the equations above are required to estimate the number of genuine driver substitutions. This is true even if a driver gene was under positive selection in every patient, as long as the extent of negative selection is negligible.

##### 5.2.1. Pentanucleotide model and removal of polymorphic sites

We have shown that using an inadequate substitution model can lead to substantial biases in the estimation of dN/dS (Supplementary Text S6, Fig. S1A). Many applications of dN/dS do not require a very high accuracy, since true biological deviations from neutrality are often far larger than the biases caused by the substitution model. For example, identification of genes under positive selection in small datasets is often unaffected by the substitution model since dN/dS ratios of genuine driver genes can take very high values (see, for example, Fig. 4E).

The estimation of the number of driver substitutions per tumor, however, requires accurate quantification of dN/dS ratios, since these ratios are often very close to 1. For example, the genome-wide dN/dS value for all non-synonymous substitutions in the pancancer dataset used in Fig. 4B is 1.059 (CI95%: 1.052, 1.065). Given the proximity to 1, misestimating this value by a few percent would have a considerable impact on the estimates of the number of driver substitutions per tumor.

To minimize the risk of systematic biases in the estimation of genome-wide dN/dS values and the average number of driver substitutions per tumor shown in Fig. 4, we took two additional precautions. First, we used a pentanucleotide context-dependent substitution model (3,072 rate parameters) instead of the trinucleotide model (192 rate parameters). Second, since somatic mutations called by our pipeline were filtered against an unmatched normal panel, common polymorphic sites in the human population (which are enriched in synonymous mutations) will be depleted of somatic mutations, which could lead to a very small upward bias in dN/dS. To entirely avoid this possible bias, all sites in the unmatched normal panel were excluded from the calculation of the numbers of synonymous and non-synonymous sites (*L*_i_) per gene.

###### Pentanucleotide model

Comparing the numbers of driver substitutions per tumor for each cancer type estimated under the trinucleotide and pentanucleotide substitution models reveals a very good agreement across all cancer types, with the exception of melanoma (Fig. S1E). This suggests that the trinucleotide model already captures the relevant context-dependent mutational biases in the data for the purpose of the estimation of the number of driver substitutions across tumor types.

The only exception is the melanoma dataset, in which the trinucleotide model estimates a significantly lower dN/dS value than the pentanucleotide model, leading to a very different estimate of the number of driver mutations per tumor. Careful examination suggests that this is due to ultraviolet-induced C>T substitutions, which show context-dependent effects extending beyond the trinucleotide context (Pleasance et al., 2010a). In fact, excluding C>T mutations from the dN/dS calculation in melanoma shows that the downward bias of dN/dS in melanoma under the trinucleotide model is largely exclusive to C>T substitutions.

To avoid these biases, all analyses in Fig. 4A-C, which require accurate dN/dS values, were carried out using the pentanucleotide substitution model.

###### List of 369 known cancer genes

To quantify the fraction of non-synonymous substitutions observed in known driver genes that are genuine driver mutations, we used a list of 369 high-confidence driver genes (Fig. 4A). This list was compiled by merging the list of 174 *COSMIC classic genes* from version 73 of the COSMIC database (Forbes et al., 2015), the list of 219 significantly mutated genes reported by Lawrence *et al*. (Lawrence et al., 2014) and the list of 204 genes identified as significantly mutated by the present study.

##### 5.3. Other mutation types: estimating the density of driver indels and synonymous mutations

Our estimates of the number of driver mutations per tumor using dN/dS are restricted to non-synonymous coding substitutions, including missense, nonsense and essential splice site substitutions. To obtain approximate estimates of the relative contribution of indels and synonymous substitutions to the number of driver mutations, we used a different approach not based on dN/dS. Briefly, the expected neutral rate of indels and synonymous substitutions on a collection of driver genes was estimated from their frequency on putative passenger genes, and this number was used to estimate the excess of these mutations observed in driver genes. This approach is conceptually analogous to the one used in (Supek et al., 2014) to estimate the frequency of synonymous driver mutations in cancer.

To estimate a background model for the *neutral* frequency of synonymous substitutions and indels we first excluded the long list of 987 putative cancer genes described in Supplementary Methods S4.4. We then used two separate negative binomial regression models for synonymous substitutions and indels. For synonymous mutations we used as an offset the expected rate of synonymous substitutions per gene under the full trinucleotide model. Unlike the approach used in (Supek et al., 2014), this entirely avoids the confounding effect of variable sequence composition across genes and trinucleotide context-dependent mutational biases. For indels we used the gene length as an offset. For both synonymous substitutions and indels, we used the 20 covariates described in Supplementary Methods S1.3, to account for the regional variation of mutation rates across the genome. These models were then applied to the list of 369 high-confidence cancer genes to estimate the number of passenger indels and synonymous mutations expected to accumulate neutrally in these genes. This enables the calculation of observed/expected ratios for synonymous substitutions and indels in known cancer genes, and, analogously to using dN/dS ratios, the estimation of the fraction of these mutations that are genuine drivers and their absolute contribution to the number of driver mutations per tumor. Confidence intervals for these estimates were obtained by bootstrapping the number of mutations observed per gene.

To evaluate the reliability of this approach, we also applied it to missense, nonsense and essential splice site substitutions and compared the estimated number of driver mutations per tumor in known cancer genes to those obtained using dN/dS. As shown in Fig. 4D, the estimates obtained from these two very different approaches are very consistent.

###### 5.3.1. Identification of cancer genes with a higher frequency of synonymous mutations than expected

Although the vast majority of synonymous mutations observed in cancer genomes are passenger mutations and accumulate largely neutrally, our analysis and a previous study (Supek et al., 2014) suggest that a small number of them can act as driver mutations. We can use the negative binomial background model for synonymous substitutions described above to identify genes with an unexpectedly high density of synonymous mutations, in a similar way in which we identify genes recurrently mutated by indel drivers (Supplementary Methods S1.5).

Running this analysis on the list of 369 cancer genes reveals that only *TP53* (q-value=6.0e-6) and *CDKN2A* (q-value=0.00058) have a convincing and statistically-significant higher than expected number of synonymous substitutions (q-value<0.01). Close inspection of the mutations in *CDKN2A* revealed that the recurrent synonymous mutations observed are indeed truncating mutations affecting a different transcript of the gene, with a different reading frame, and so *CDKN2A* is not genuinely recurrently affected by synonymous driver mutations.

*TP53* has been previously reported to be the target of synonymous driver mutations (Supek et al., 2014), which affect the correct splicing of the transcript, and our analysis entirely supports this conclusion. The observed/expected ratio of synonymous substitutions in *TP53* is very high (∼6.8), which suggests that a majority of the synonymous mutations observed in *TP53* in our cohort of 24 cancer types are likely genuine driver mutations. In fact, in our cohort, over half of the synonymous substitutions observed in *TP53* affect the same site T125T (21 out of 39 synonymous substitutions in *TP53*), a recurrent synonymous hotspot known to lead to aberrant splicing (Supek et al., 2014). Hence, this single synonymous hotspot accounts for the majority of synonymous driver substitutions in *TP53*, although other synonymous mutations in *TP53* are also likely drivers.

A previous study identified a number of oncogenes with a higher density of synonymous mutations than expected by chance (Supek et al., 2014) and argued that these could be driver mutations affecting splicing. Among these genes, the study highlighted 11 oncogenes with a particularly high density of synonymous substitutions: *PDGFRA*, *EGFR*, *KDR*, *NTRK1*, *IL7R*, *TSHR*, *ELN*, *JAK3*, *ITK*, *GATA1* and *RUNX1T1*. This contrasts with our analysis, which only identified *TP53* as having a statistically-significant higher rate of synonymous mutations despite using a dataset with nearly twice as many samples as the dataset used in the previous study. An important difference between our analysis and that in the previous report is that our negative binomial model uses overdispersion to quantify the uncertainty in the estimated mutation rate for a gene. This makes our model more conservative, but also more robust against false positives caused by the neutral variation of the mutation rate across genes. We also control for trinucleotide sequence composition and trinucleotide mutation rates as well as 20 epigenomic covariates in the estimation of the background mutation rate per gene. Interestingly, even though both studies are based on TCGA samples and, in fact, share a large number of samples, only 4 of the 11 oncogenes highlighted in the previous study as having a high rate of synonymous mutations have observed/expected ratios of synonymous substitutions >1.5x according to our model and none are considered significant under the negative binomial model (q-value>0.5).

Overall, consistently with previous reports, our analyses suggest that certain synonymous mutations can indeed act as cancer driver mutations, of which the T125T hotspot mutation in *TP53* is probably the most striking example. However, there is little evidence that this is a general and frequent mechanism. Our analyses suggest that synonymous mutations contribute a small fraction (<5%) of all driver mutations seen in cancer genomes (Fig. 4D).

### Supplementary Text

#### 6. Simplistic substitution models lead to biased dN/dS ratios and false inference of selection

Traditional implementations of dN/dS have typically used simplistic substitution models. The classic implementation of dN/dS by Nei and Gojobori (Nei and Gojobori, 1986), for example, uses a substitution model in which all substitutions are equally likely (F81 substitution model). More sophisticated likelihood implementations of dN/dS, such as the widely used implementation in the *PAML* software package, typically use a simple substitution model with a different rate for all transitions (C<>T and G<>A changes) and all transversions (C<>A, C<>G, G<>C and G<>T changes) (HKY85 substitution model) (Goldman and Yang, 1994; Yang, 2007). A more complex substitution model, frequently used in molecular evolution but more rarely in dN/dS analyses, is the GTR (General Time Reversible) model, which has 6 mutation classes, one for each of the 6 possible reversible base changes (A<>C, A<>G, A<>T, C<>G, C<>T, G<>T).

In reality, the substitution rate often varies markedly depending on the exact nucleotide change and on the bases upstream and downstream of a base. This is particularly well understood in cancer, from the study of mutational signatures (Alexandrov et al., 2013). The use of simplistic mutation models is known to lead to biases in dN/dS estimates (Yang and Nielsen, 2000). While these biases may be of lesser importance in the presence of overwhelming negative or positive selection, they can have important implications when dN/dS ratios are close to 1, as is often the case in somatic evolution.

Fig. S1A reveals how simplistic substitution models lead to systematic under or overestimation of dN/dS ratios and to wrong inference of selection. To generate this figure, the average trinucleotide substitution rates (192 parameters) were estimated in three different cohorts of samples, which are dominated by different mutational processes: pancancer (dominated by C>T mutations at CpG sites), melanoma (dominated by the UV-signature of C>T mutations at cytosines with a pyrimidine upstream) and lung adenocarcinoma (dominated by G>T mutations generated by tobacco smoking) (Alexandrov et al., 2013). Using the trinucleotide rates observed in each of these datasets, and the trinucleotide frequencies of the human exome, we simulated 100 datasets with 10,000 random coding substitutions per dataset. The correct dN/dS ratio in these simulations is 1, since the mutations were simulated entirely randomly, without selection. Fig. S1A shows how estimated dN/dS ratios under different simplistic substitution models systematically deviate from the correct value of 1. In fact, these biases are large enough to suggest considerable negative and positive selection when using simplistic models.

These biases have important implications. For example, a study applying dN/dS to somatic mutations from breast cancer genomes used a Nei-Gojobori implementation of dN/dS (F81 substitution model), obtaining a global dN/dS∼0.82 (Ostrow et al., 2014). This led the authors to conclude that weak negative selection operates in cancer somatic mutations, when in reality this dN/dS ratio is a consequence of the downward bias in dN/dS under the Nei-Gojobori model (Fig. S1A).

#### 7. Trinucleotide vs pentanucleotide substitution models

The use of a full trinucleotide model comprehensively accounts for the majority of known context-dependent mutational biases. Previous studies suggest that context dependent effects beyond three nucleotides are relatively small (Alexandrov et al., 2013).

To evaluate the impact on dN/dS of context-dependent effects extending beyond one base up-and downstream, we compared whole-genome estimates of dN/dS across cancer types under the full trinucleotide (192 rate parameters) and a full pentanucleotide model (3,072 rate parameters). Fig. S1D,E reveals that the addition of context-dependent effects beyond three nucleotides does not have a significant impact on genome-wide dN/dS ratios in any cancer type, with the exception of melanoma. As discussed in Supplementary Methods S5.2, this is due to UV-induced C>T mutations showing context-dependent effects extending beyond the trinucleotide level (Pleasance et al., 2010a).

With the exception of melanoma, in which the dominant mutation processes lead to a slight downward bias in dN/dS under the trinucleotide model, Fig. S1D,E evidence that the trinucleotide model captures most of the relevant context-dependent effects required for a very accurate estimation of dN/dS.

#### 8. Impact of germline SNP contamination or SNP over-filtering

As shown in Fig. 1A, coding germline SNPs are heavily enriched in synonymous mutations as a result of purifying selection on germline mutations during human evolution (dN/dS ratios for missense and truncating substitutions are 0.38 and 0.08, respectively). Identification of somatic mutations in cancer genomes requires careful removal of germline polymorphisms by sequencing a matched normal sample in addition to a tumor sample from each patient. Given the action of negative selection on germline SNPs, incomplete removal of SNPs from catalogs of somatic mutations can introduce a false signal of negative selection. To protect against germline SNP contamination, some pipelines systematically remove putative somatic mutation overlapping polymorphic sites in humans in addition to using a matched normal sample. However, since polymorphic sites are enriched in synonymous sites, such filtering strategy can lead to over-filtering of genuine somatic mutations, with a bias against synonymous sites.

Fig. S1B,C show how both germline SNP contamination and over-filtering of SNP sites can have a considerable impact on global dN/dS ratios, resulting in signals of negative and positive selection, respectively. To generate this figure, we first simulated ten neutral datasets of somatic mutations by randomization of existing cancer genomic datasets (see Supplementary Text S9). To these neutral datasets, we added 5% or 10% of randomly selected germline SNPs (Fig. 1) or we subtracted any mutation overlapping known polymorphic sites using the dbSNP database.

Interestingly, this analysis confirms that global dN/dS ratios detect a very clear signal of negative selection even when only 5% of all mutations are germline SNPs. This further emphasizes the remarkable lack of negative selection reported in Fig. 3, and in particular in Fig. 3G after comprehensively removing known cancer driver genes.

SNP contamination and SNP over-filtering are likely to affect TCGA public somatic mutation calls from different datasets to different extents. This was apparent when we calculated global dN/dS ratios using the somatic mutation calls publicly released by TCGA. For example, the *COAD*, *READ* and *KICH* datasets showed significantly lower dN/dS ratios than expected: *COAD* = 0.92 (CI95%: 0.91, 0.94), *READ* = 0.91 (CI95%: 0.87, 0.95) and *KICH* = 0.94 (CI95%: 0.89, 1.00), suggesting the presence of SNP contamination in these datasets. To determine whether these low dN/dS ratios are truly caused by SNP contamination of the public catalogs of somatic mutations, we calculated the fraction of mutation calls overlapping common germline SNP sites (dbSNP database build 146). This revealed that these three datasets have a much higher fraction of somatic calls overlapping common dbSNP sites than other datasets, with 11.0%, 15.6% and 12.2% of all somatic mutation calls from TCGA overlapping known SNP sites (Fig. S1C). In contrast, the median percentage of overlapping calls in all other cancer types from TCGA is 1.7% (range: 0.66-3.3%).

Studies searching for evidence of negative selection based on public mutation calls from TCGA are likely to be affected by the confounding effects of SNP contamination and potentially SNP over-filtering. Having control over the strategy for SNP filtering was the main motivation for uniformly re-calling somatic mutations across TCGA datasets in the present study. In order to minimize the risk of germline SNP contamination, we required a minimum coverage of 10x in the matched normal sample of a putatively mutated site. To entirely avoid any risk of over-filtering of SNP sites that may introduce an upward systematic bias to dN/dS, we did not perform dbSNP filtering and all sites masked out by our unmatched normal panel were excluded from the calculation of available sites (*L*) in dN/dS. Reassuringly, *CaVEMan* somatic mutation calls for all TCGA datasets showed the expected low overlap with common dbSNP sites, with a median of 1.8% (range: 1.1-3.2%), a figure consistent with the expectation from neutral simulations.

#### 9. Performance of different dN/dS models for driver discovery

Previous studies have highlighted the importance of adequately modeling the variation of mutation rates along the genome to identify driver (positively selected) genes with good specificity and sensitivity. Particularly, Lawrence *et al*. (Lawrence et al., 2013) showed how models that do not account for the regional variation of the mutation rate along the genome can yield very long lists of false positives.

To evaluate the specificity of different methods in the presence of realistic levels of mutation rate variation along the genome, we can use realistic neutral simulations of somatic mutations. In line with ongoing international benchmarking efforts of driver discovery methods, we generated simulated neutral datasets by local randomization of somatic mutations from real whole-genome sequencing studies. Using data from 107 melanoma whole-genomes from ICGC, we first filtered out coding mutations from a panel of known driver genes, to minimize the presence of driver mutations, and then reassigned each mutation to a randomly selected position with an identical trinucleotide context within 50kb of its original position. This randomization procedure results on a neutral dataset that retains the same variation of mutation rates and mutational signatures across patients and across regions of the genome.

In a neutral dataset, robust methods for driver discovery with good specificity should not yield any significant hit. This can be formally evaluated by performing false discovery rate correction and by plotting the vector of *P*-values under the null model (neutral simulation) in a QQ-plot. The QQ-plot in Fig. S1F reveals that the *uniform rate dN/dS model* yields a large number of false positives in the neutral simulation described above, as expected in the presence of large neutral variation of the mutation rate along the genome (Lawrence et al., 2013). In contrast, both the *variable rate dN/dS model* (which estimates the local mutation rate from the synonymous substitutions in each gene) and *dNdScv* (which uses the regression framework described in section S1.3 in addition to local synonymous substitutions) have perfect specificity under the challenging conditions of the simulation above (Fig. S1F). This result is representative of simulations performed under a variety of assumptions and starting datasets, even using simulated datasets with thousands of samples. The specificity of *dNdScv* has also been demonstrated by an international benchmarking exercise as part of the Pancancer Analysis of Whole-Genomes Consortium (PCAWG-ICGC) [*manuscript in preparation*].

Although both the *variable rate dN/dS model* (which we used in (Wong et al., 2014)) and *dNdScv* have good specificity under challenging conditions, they differ dramatically in terms of their sensitivity. This is shown in Fig. S1G, which depicts the number of significant genes identified by both methods across the TCGA datasets analyzed in this study. While the *variable rate dN/dS model* can detect a substantial number of positively selected genes in large datasets (Wong et al., 2014), *dNdScv* has higher sensitivity across datasets of any size, both when analyzing substitutions alone or when combining substitutions and indels. *dNdScv* has also been found to have similar or higher sensitivity than all other driver discovery methods benchmarked in the Pancancer Analysis of Whole-Genomes Consortium (PCAWG-ICGC) [*manuscript in preparation*], including *MutSgCV* (Lawrence et al., 2014) and *oncodriveFML* (Mularoni et al., 2016).

#### 10. Factors contributing to the weakness of negative selection in somatic evolution

Negative selection on somatic mutations during cancer evolution has been long anticipated (Beckman and Loeb, 2005; McFarland et al., 2014; Nowell, 1976; Stratton et al., 2009). However, several authors have also predicted that, while present, negative selection should be weaker in somatic evolution owing to a number of factors (*e.g.* (McFarland et al., 2013; Morley, 1995)). Some relevant factors are listed below:

1. *Diploidy*: Having two (or more) copies of every gene is a major buffer against negative selection (Morley, 1995), as shown in Fig. 3G. This is different for germline mutations, since, owing to sexual recombination, selection acts on recessive deleterious alleles by purging homozygous individuals. The loss of a single copy of a gene can still have deleterious effects owing to haploinsufficiency, but selection against such alleles will still be expected to be weaker in the absence of sexual recombination.
2. *Large fraction of dispensable genes in somatic cells*: For any given somatic lineage a large number of genes are likely to be dispensable (Morley, 1995), as shown by Fig. 3G. This is different for germline mutations, which will be exposed to selection even if they only manifest as deleterious in certain tissues, in certain conditions or in certain stages of development, for example. Also, it has been argued that somatic cells defective in certain functions can still prosper within a tissue by exploiting the effort of wild-type cells around them (Morley, 1995).
3. *Frequent hitchhiking with drivers*: Very weakly deleterious alleles (*e.g.* alleles reducing the survival probability of a cell per year by a few percent) require time to be effectively purged from a population. The random occurrence of a very advantageous mutation, such as a cancer driver mutation, in a cell carrying weakly deleterious mutations can offset their fitness effects and lead to the fixation of deleterious mutations in a cancer. Frequent rounds of clonal expansions and hitchhiking with potent driver mutations is expected to significantly reduce the efficiency of negative selection to remove deleterious variants (McFarland et al., 2013; McFarland et al., 2014).
4. *Muller’s ratchet*: Somatic evolution is effectively asexual and so deleterious mutations will be expected to accumulate in somatic cells even in the absence of hitchhiking with driver mutations (McFarland et al., 2013; McFarland et al., 2014). Muller’s ratchet is expected to be stronger the higher the mutation rate per division, which predicts that negative selection will be even weaker in hypermutator samples.

All of the factors above, among others, are likely to contribute to the observed weakness of negative selection during cancer evolution. Nevertheless, the extreme weakness of negative selection observed in cancer genomes remains surprising, extending to mutations with dominant phenotypes, including coding mutations predicted to create neoantigens, as well as truncating mutations in highly expressed genes and haploid regions of the genome.

**Figure S1.**
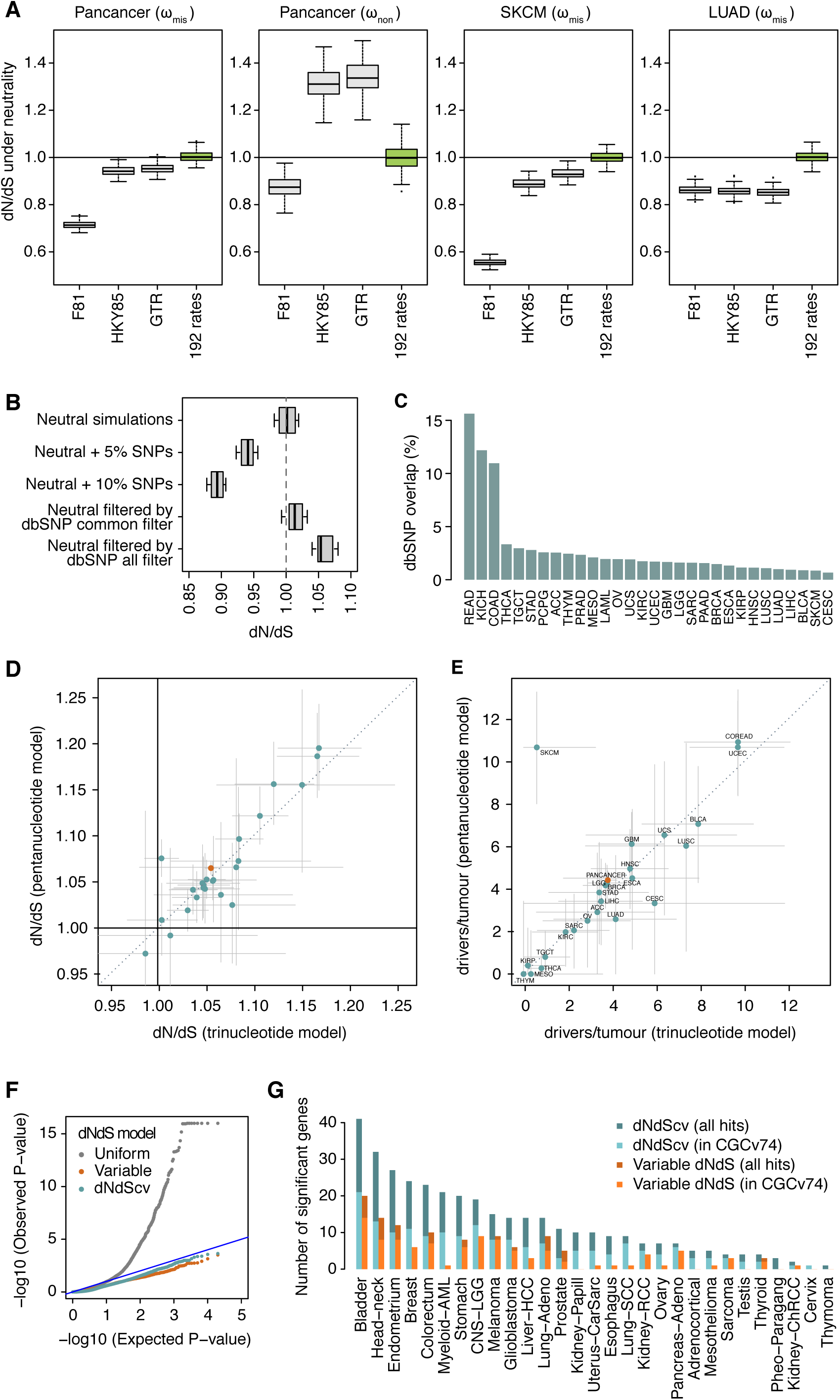
[*Related to Figures 1-4*] Impact of different confounding factors on analyses of selection, including simplistic substitution models, SNP contamination, SNP filtering and inadequate background models of the variation of the mutation rate. (**A**) Impact of simplistic mutation models on the accuracy of dN/dS in different scenarios. Each boxplot represents the dN/dS ratios estimated from 100 neutral simulations of 10,000 random coding substitutions. To exemplify the impact on dN/dS of different mutational spectra, we simulated neutral datasets using the trinucleotide spectra observed in the three different cohorts of samples (pancancer, melanoma and lung adenocarcinoma). Different panels depict dN/dS ratios for missense (ω_mis_) or nonsense (ω_non_) mutations. (**B**) Simulations of the impact on dN/dS of germline SNP contamination and SNP over-filtering in catalogs of somatic mutations. 10 neutral datasets were generated by local randomization of 607 cancer whole-genomes (Alexandrov et al., 2013), as described in Supplementary Text S9. Datasets with varying degrees of germline SNP contamination were simulated by adding 5% or 10% of germline common SNPs (minor allele frequency >=5%) from 1000 genomes phase 3 (Auton et al., 2015) to the neutral simulations. Datasets with varying levels of SNP over-filtering were simulated by removing any mutation from the neutral datasets that overlapped a polymorphic site in dbSNP build 146 (either using common sites or all sites) (Sherry et al., 2001). (**C**) Percentage of mutations from the public TCGA catalogs of somatic calls that overlap a common dbSNP site. Based on simulations, an overlap of 1-3% might be expected depending on the dominant mutational signatures present in a dataset (Supplementary Text S8), but several public TCGA catalogs show a much higher overlap suggesting extensive germline SNP contamination. As predicted from Fig. S1B, this leads to an artefactual signal of negative selection in these datasets (Supplementary Text S8). (**D**) Consistency between genome-wide dN/dS estimates using the trinucleotide and pentanucleotide substitution models across cancer types. Green dots represent genome-wide dN/dS estimates for each cancer type separately, and the orange dot depicts the pancancer estimates (using the 24 cancer types with *CaVEMan* mutation calls). (**E**) Corresponding estimates of the average number of driver coding substitutions per tumor, calculated as described in Supplementary Methods S5.2. For the purpose of estimating the excess of mutations from dN/dS ratios, dN/dS values below 1 are set to 1. Error bars depict confidence intervals 95%. (**F, G**) Evaluation of the relative performance of the three different dN/dS models for the detection of positive selection at gene level (driver gene discovery). (**F**) QQ-plots for the different dN/dS models on a neutral dataset obtained by randomization of 107 melanoma whole-genomes from ICGC (Supplementary Text S9). The *uniform rate dN/dS model* displays a great inflation of low *P*-values, leading to a large number of false positives after multiple testing correction (368 genes with q-value<0.05), and should be generally avoided. In contrast, both the *variable rate dN/dS model* and *dNdScv* behave as expected for a neutral dataset, yielding no significant hits after multiple testing correction. (**G**) Sensitivity of *dNdScv* and of the *variable rate dN/dS model*. The bar plot depicts the number of significant genes (q-value<0.05) identified by both methods in the 29 TCGA datasets. Bars colored in a lighter shade show the number of significant genes that are present in the Cancer Gene Census version 74 (Forbes et al., 2015). *dNdScv* shows good specificity and sensitivity under all tested conditions.

**Figure S2.**
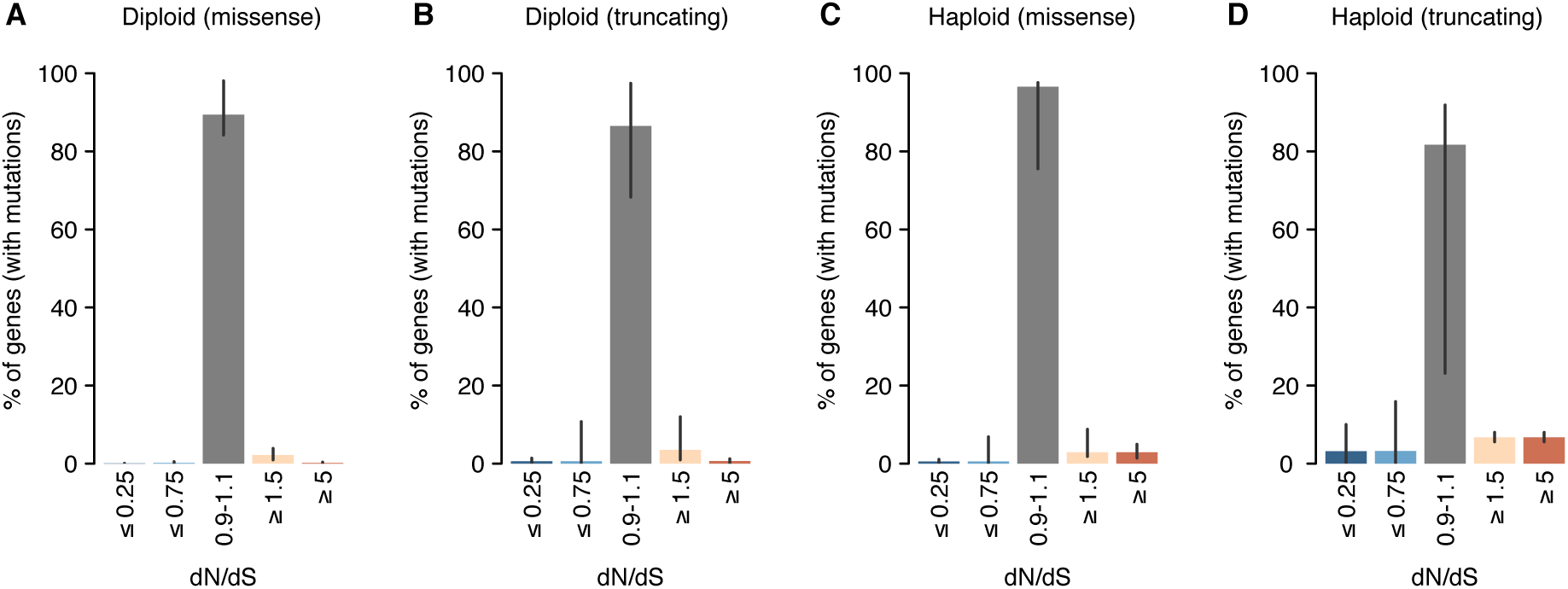
[*Related to Figure 3*] dN/dS distributions inferred for different mutation types and copy number states. These distributions, obtained as described for Fig. 3C, represent the percentage of genes estimated to be under a certain selection regime. The four distributions correspond to: missense (**A**) and truncating (**B**) substitutions in regions without loss of heterozygosity, and missense and truncating substitutions in haploid regions (**C** and **D**, respectively). Note that Fig. S2A is an extension of Fig. 3C, with an added middle bar for genes with dN/dS very close to 1 (0.9-1.1), which can be considered to evolve largely neutrally. Only samples with *CaVEMan* mutation calls, excluding melanoma samples, were considered for this analysis for the reasons explained in Supplementary Methods S4.1. For each figure, all mutations with the appropriate ploidy were included in the analysis and only genes with at least one mutation (either synonymous or non-synonymous) participate in the fitting of dN/dS distributions (Supplementary Methods S4.2.2). Hence, the percentages of genes shown in the y-axes are relative to the total number of genes with at least one mutation in regions with the ploidy considered in each figure. Error bars depict 95% confidence intervals.

**Figure S3.**
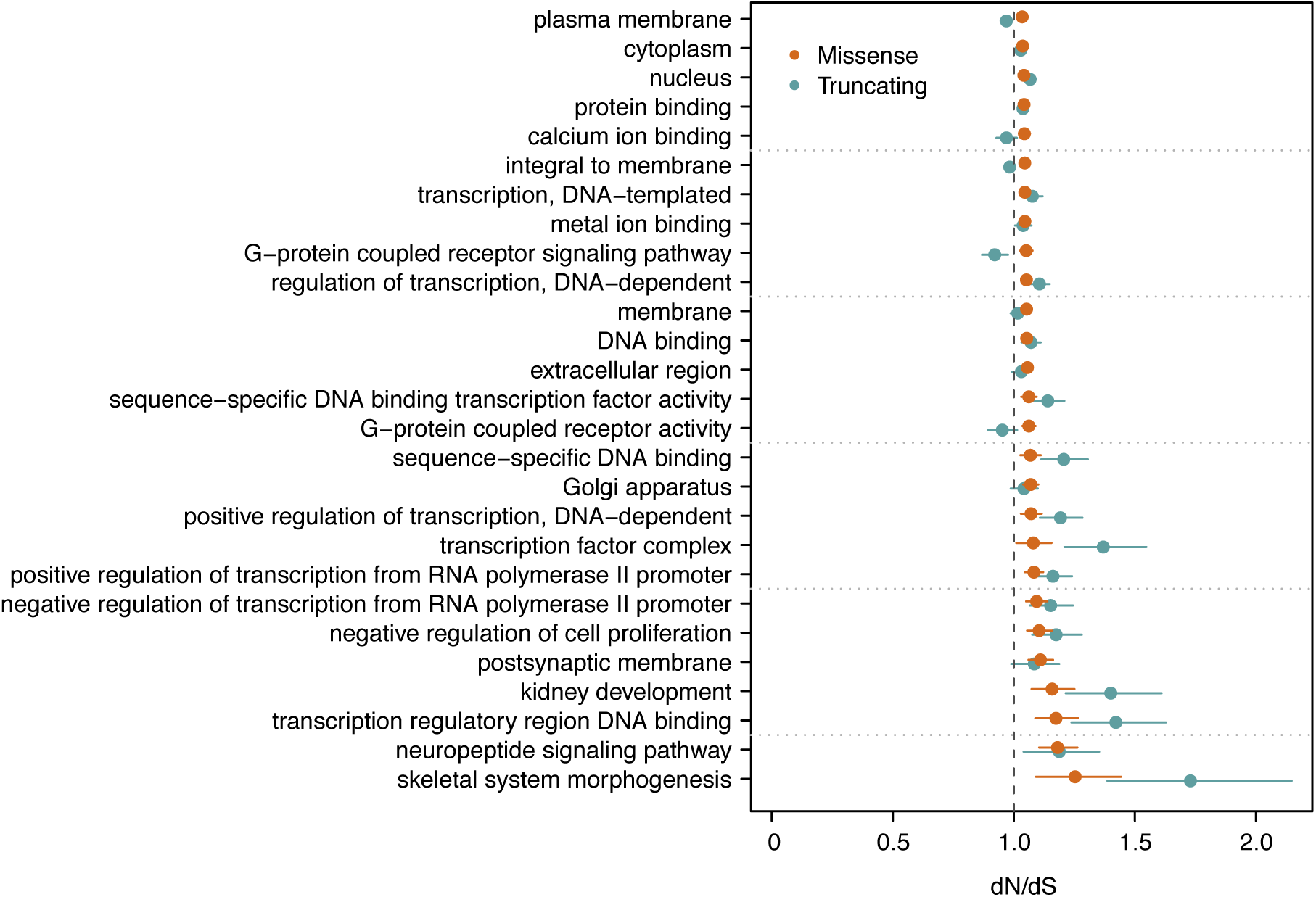
[*Related to Figure 3*] Gene ontology groups deviating significantly from neutrality after removing known cancer genes. See Supplementary Methods S4.4 for a detailed description of this analysis. 27 gene ontology classes are found to be under significant positive selection after comprehensively removing 987 known putative cancer genes. This suggests the presence of undiscovered cancer genes in these functional groups. No gene ontology class was found to be under significant negative selection. Error bars depict 95% confidence intervals.

**Figure S4.**
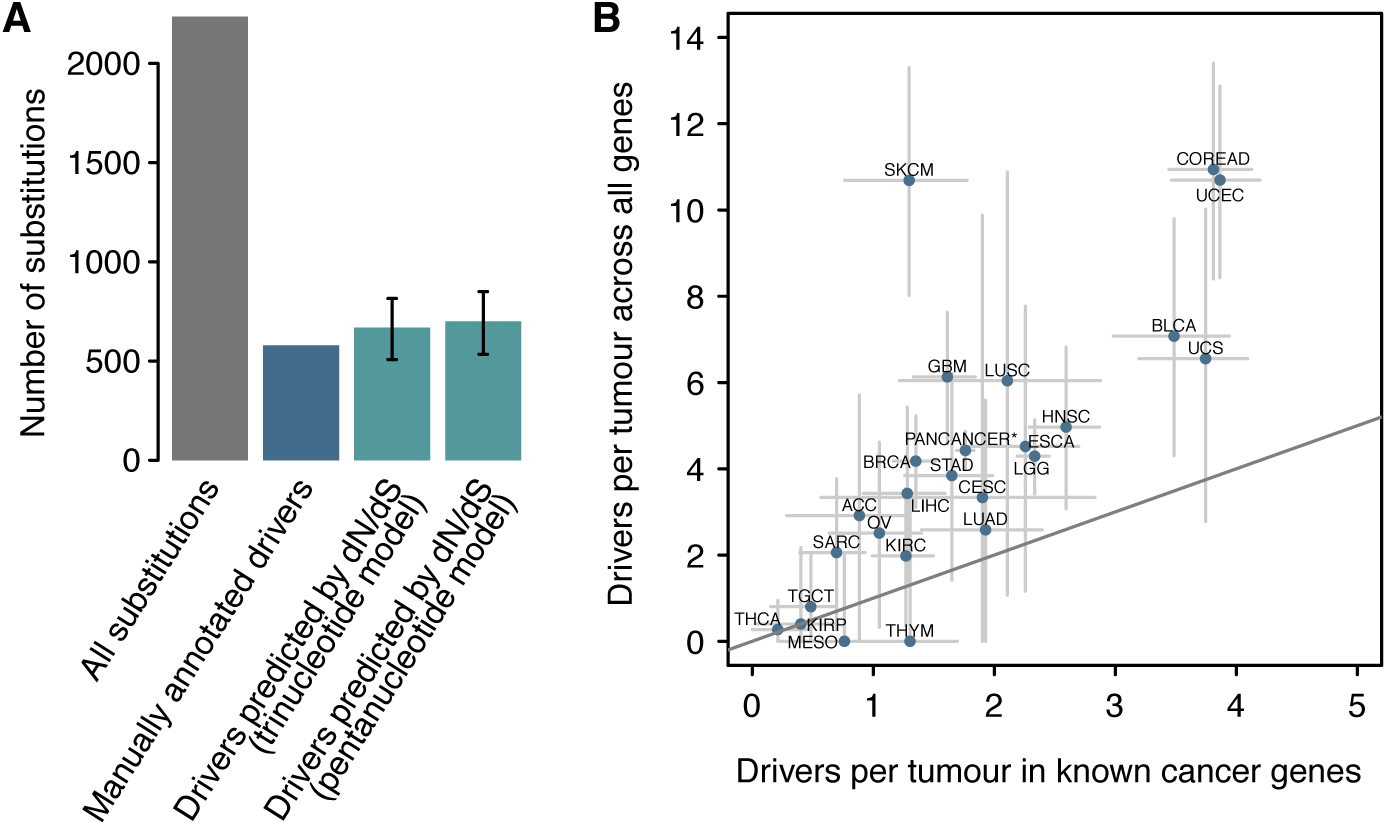
[*Related to Figure 4*] Supplementary analyses on the number of coding driver substitutions per tumor. (**A**) Comparison of the number of coding driver substitutions estimated by dN/dS and the number estimated by manual annotation of driver mutations across 560 breast cancers. The figure depicts the total number of coding substitutions (grey bar) and the estimated number of driver substitutions in a list of 723 putative cancer genes across 560 breast cancer whole-genomes. A total of 2,786 coding substitutions are found in these genes across the 560 patients (data from (Nik-Zainal et al., 2016)). Of these, 579 were annotated as likely driver mutations by a careful and conservative manual curation in the original publication (Nik-Zainal et al., 2016) (blue bar). Using the trinucleotide dN/dS model on this dataset, restricted to these 723 genes, yielded a global dN/dS for all non-synonymous substitutions of 1.42 (CI95%: 1.29, 1.58). Reassuringly, this led to an estimated number of drivers consistent with the manual annotation: 668.9 (CI95%: 507.5, 815.3). Error bars depict confidence intervals 95%. (**B**) Scatter plot of the estimated average number of coding driver substitutions per tumor in 369 known cancer genes and in all genes of the genome. This is a scatter plot representation of the bottom panels of Fig. 4A,B, to emphasize the extent of coding driver substitutions occurring outside of the list of 369 cancer genes. Error bars depict confidence intervals 95%. Note that the two cancer types whose estimates appear under the diagonal (mesothelioma –MESO-and thymoma –THYM-) have confidence intervals extending above the diagonal, as expected.

